# PKD2L1 channels segregated to the apical compartment are the dual-mode pH sensor in cerebrospinal fluid-contacting neurons

**DOI:** 10.1101/2025.08.24.671961

**Authors:** Magdalena Vitar, Daniel Prieto, Stavros Malas, Raúl E. Russo, Federico F. Trigo

## Abstract

Cerebrospinal fluid contacting neurons (CSFcNs) are GABAergic cells that surround the central canal (cc) of the spinal cord. Their soma is located sub-ependymally and they have a dendritic-like process that ends as a bulb (the so-called “apical process”; ApPr) inside the cc. It remains unclear how this unique anatomical organization, with the soma and the ApPr located in different extracellular environments, relates to their function as a multimodal sensor of cerebrospinal fluid (CSF) composition. One of the main physiological features of CSFcNs is a prominent spontaneous electrical activity mediated by PKD2L1 channels, a non-selective cation channel of the TRP family. PKD2L1 channels have a high single-channel conductance (around 200 pS) and can be modulated by protons and mechanical forces. In this work we investigate PKD2L1 channel sensitivity to pH and its effects on CSFcNs excitability. We demonstrate that PKD2L1 spontaneous activity generates not only phasic inward currents, but also a sustained current, both of which are modulated bidirectionally by pH with a high sensitivity around physiological values. By combining electrophysiology (direct recordings from intact and isolated ApPrs) with optical methods (laser-photolysis of protons) we further show that functional PKD2L1 channels are specifically localized in the ApPr. The spatial segregation of PKD2L1 channels, along with their biophysical properties (high single-channel conductance and pH sensitivity) and the ApPr’s unique membrane properties (very high input resistance) renders CSFcN excitability exquisitely sensitive to PKD2L1 modulation. Altogether, our findings illustrate how the ApPr’s properties are finely tuned to support its sensory role.

## Introduction

Sensory information originates from the stimulation of specific receptors in different parts of the body. Located at the interface between the spinal cord central canal (cc) and the spinal cord parenchyma, cerebrospinal fluid-contacting neurons (CSFcNs) are part of a sensory system that provides information about the internal environment, specifically the composition of cerebrospinal fluid (CSF), and are thus considered part of the interoceptive system^1,2^.

Sensory cells transduce sensory stimuli into electrical activity and typically have distinctive morphological specializations. CSFcNs are no exception, with their somas located in the subependymal layer from which a short, thick dendrite arises, ending in a bulbous structure located within the cc^3,4^ known as the “apical process” (ApPr). The axon of CSFcNs, on the other hand, projects rostrally trough the ipsilateral ventral spinal cord, and contacts other CSFcNs and neurons of spinal central pattern generators, including motorneurons and premotor excitatory neurons^5–7^. By providing sensory information about CSF composition and the mechanical forces acting near the cc, CSFcNs participate in the control of posture and locomotion, both in lower vertebrates^1,5,8–10^ and in rodents^6,7^.

CSFcNs are established chemoreceptors. Huang et al. were the first to report that CSFcNs in the mouse spinal cord: i. express the PKD2L1 channel, a cation permeable channel of the TRP (Transient Receptor Potential) family that has a high Ca^++^ permeability^11^; ii. respond to acid with an increase in firing rate^12^. Subsequent studies examined the pH sensitivity of CSFcNs in the rat^3^, lamprey^10,13^ and mouse spinal cord^14,15^. These collective findings suggested that the response of CSFcNs to acidification depends on the activation of an acid sensing ion channel (ASIC) (which produce a rapidly desensitizing inward current) and the response to alkalinization depends on the activation of PKD2L1 channels.

Despite previous work, the precise mechanisms and subcellular localization of the channels responsible for the pH response in CSFcNs remain unknown. This gap is partly due to two main factors. First, earlier investigations relied on bath application or pressure ejection of solutions with varying pH, which lack the spatial and temporal precision required to resolve localized pH sensitivity in specific cellular compartments. Second, there are technical challenges associated with recording from a small neuronal compartment using electrophysiological approaches. Here, we overcome these technical barriers with a combination of whole-cell and outside-out recordings directly from the ApPr, along with laser photolysis of protons and immunohistochemistry. We reveal that PKD2L1 generates not only phasic but also a novel sustained current that critically shapes the membrane potential of CSFcNs. These currents exhibit high pH sensitivity near the physiological range. Importantly, we demonstrate that functional PKD2L1 channels are predominantly localized to the ApPr, whose high input resistance and pH sensitivity make it a highly specialized sensory hub for the integration of chemical signals.

## Results

### PKD2L1-mediated spontaneous activity in CSFcNs from Gata3^eGFP^ mice

In this work we used the Gata3^eGFP16^ transgenic mice, where CSFcNs express eGFP under a GATA3 regulatory element. GATA3 is a transcription factor expressed in the spinal cord both by CSFcNs^17^ and V2b interneurons^16^. **Figure 1A** shows sagital (**Aa**) and transverse (**Ab**) slices of Gata3^eGFP^ animals where the typical morphology and location (around the cc) of CSFcNs can be readily appreciated. This, together with the fact that V2b interneurons are located well away from the cc, allows unambiguous identification of CSFcNs. As expected, GFP+ in the Gata3^eGFP^ mice express the PKD2L1 channel showing they are indeed CSFcNs (**Figure 1Ac** and **d**). We first characterised the basal electrophysiological activity of CSFcNs from Gata3^eGFP^ mice in voltage-clamp, as these neurons in the Gata3^eGFP^ transgenic have not yet been characterized. CSFcNs were recorded with a KGluconate-based intracellular solution (IS) at near physiological temperature (34 ± 1 °C) with the whole-cell configuration of the patch-clamp technique from either the soma or the ApPr; neurons were clamped at -60 mV. In these conditions, spontaneous single-channel events were recorded, confirming previous results (in mice^14,15^, lamprey^10^ and zebrafish^18^). **Figure 1Ba** shows a 1 sec period of such a recording. The dotted horizontal green lines indicate 4 levels of channel activity (c = closed; o1 = 1 channel open; o2 = 2 channels open; o3 = 3 channels open). **Figure 1Bb** shows the corresponding histogram, where the 4 peaks that define each level of channel activity are shown with arrowheads. From the analysis, the apparent probability of each state (pc, po1 and po2) was calculated. In this example, the single channel current recorded at -60 mV holding potential was

**Fig 1.**
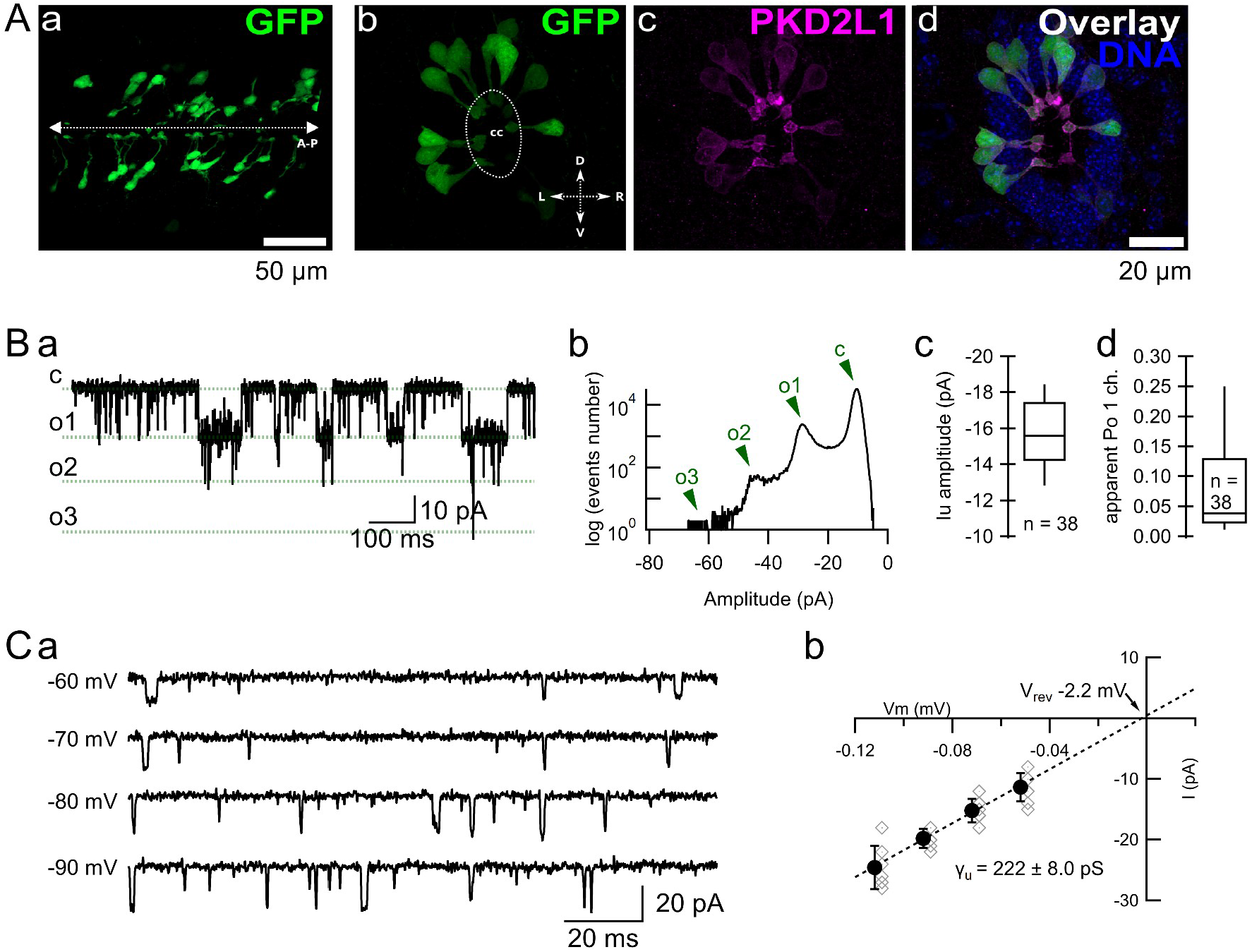
Characterization of PKD2L1 channel activity in CSFcNs from GATA3 mice. **Aa.** Confocal image of a sagital spinal cord slice (the dissection plane passes through the cc) from a Gata3^eGFP^ animal showing the distribution of CSFcNs in the anterior-posterior (A-P) direction. **Ab, c** and **d**. Confocal image of a coronal spinal cord slice from a Gata3^eGFP^ animal showing the distribution of CSFcNs (**b**, green) around the cc (dotted line) and the immunoreactivity against the PKD2L1 channel (**c**, magenta). The overlay of the 2 channels plus the DNA (blue) are shown in **d**. In **b**, D is dorsal, V is ventral, L is left and R is right. Brightness and contrast were adjusted for display purposes. **Ba.** Spontaneous activity of a CSFcNs recorded at -60 mV at 34 °C. The green, horizontal dotted lines represent the different states of the channel: c = closed; o1 = 1 channel open; o2 = 2 channels open; o3 = 3 channels open. **Bb.** Histogram from the whole-cell current shown in **a**, with the 4 peaks corresponding to c, o1 and o2 (and o3) indicated with arrowheads. **Bc.** Boxplot showing the unitary current amplitude measured from 38 different neurons (either somatic, n = 13, or ApPr, n = 25, whole-cell recordings). Middle horizontal line shows the median value (-15.6 pA), upper and lower horizontal lines the 75th and 25th percentiles, respectively, and top and low whisker the 90th and 10th percentiles, respectively. **Bd.** Boxplot showing the apparent open probability for a single channel measured from 38 different neurons (either somatic, n = 13, or ApPr, n = 25, whole-cell recordings). Middle horizontal line shows the median value (0.04), upper and lower horizontal lines the 75th and 25th percentiles, respectively, and top and low whisker the 90th and 10th percentiles, respectively. **Ca.** Spontaneous activity recorded at different holding potentials. **Cb.** IV relationship contructed from recordings performed at -52 (n = 9), -72 (n = 16), -92 (n = 14) and -102 mV (n = 7) holding potentials. Black symbols show averages ± SDs, and gray diamonds show individual values. The dotted line is a linear fit to the average data. From this fit an average unitary conductance of 222 ± 8.0 pS was calculated, with an extrapolated reversal potential of -2.2 mV. The Vm values have been corrected for a calculated liquid junction potential of -12 mV. In **B**, open diamonds correspond to individual neurons and filled circles to the mean ± SD.

-17.5 pA, and po1 was 0.12. From the spontaneous activity of different CSFcNs recorded either from the ApPr or from the soma, we measured an average single channel current amplitude of -15.9 ± 1.8 pA (**Figure 1Bc**, n = 38, with minimal and maximal values of -12.1 and -19.3, respectively) and a highly variable average apparent single channel open probability, po1, of 0.08 ± 0.08 at -60 mV (**Figure 1Bd**, n = 38, with minimal and maximal values of 0.0065 and 0.29, respectively). No difference was found for these two values between ApPr and somatic recordings (p = 0.06 for the single channel current and p = 0.53 for po1), and the results were pooled together (25 recordings from the ApPr and 13 from the soma). **Figure 1Ca** shows the single-channel activity recorded at different holding potentials from the same neuron shown in **B**. From 16 cells recorded at different holding potentials, we calculated a single channel conductance of 222 ± 8.0 pS ( **Figure 1Cb**, similar to what has been described before in CSFcNs from different species^13,14^ and for the PKD2L1 channel expressed in heterologous expression systems), with an extrapolated reversal potential very close to the expected value of 0 mV (-2.2 mV). These experiments indicate that the single channel activity recorded from adult CSFcNs from the Gata3^eGFP^ mice is very similar to previous recordings in other transgenic mice models^14,15^. The very high single channel conductance of this current and the fact that it can be blocked by dubicaine hydrochloride (see next section) indicate that this is a PKD2L1-dependent current.

### PKD2L1 channels mediate phasic and sustained currents

Dibucaine hydrochloride was described as a blocker of PKD2L1 channels in an expression system^19^. We decided to test whether dibucaine could also be used as a blocker of these channels in CSFcNs. As shown in **Figure 2Aa**, pressure-application of 100-200 µM dibucaine during 30 seconds strongly reduced the frequency of single channel openings, from 177 ± 163 to 5 ± 8 Hz (control vs dibucaine, respectively, **Figure 2Ab**, n = 12, p = 0.0005). These experiments indicate that dibucaine can be used as a pharmacological tool to study PDK2L1-dependent effects.

Apart from blocking PKD2L1 phasic activity, dibucaine had another remarkable effect. Pressure application of the drug decreased the holding current, as shown in the representative example in **Figure 2Ba** and **b**, where the holding current (HC) went from -23 to -10 pA. On average, dibucaine decreased the HC from -18.3 ± 9.5 to -10.5 ± 6.6 pA (**Figure 2Ca**, n = 13, p = 0.0002), a 44 ± 12 % reduction. Pressure application of the extracellular solution without dibucaine had no effect on either the single-channel current activity or the HC (data not shown). As expected from the effect on the HC, when the CSFcN was recorded in current-clamp (I-clamp), pressure application of dibucaine induced a marked hyperpolarization of the resting membrane potential (RMP, see **Figure 2Bc**), ffrom -66.8 ± 11.0 to -89.0 ± 14.0 mV (**Fig 2Cb**, n = 11, p = 0.001). Because dibucaine is a local anaesthetic it may act on other targets, like voltage-dependent sodium and calcium channels ^20^. To rule-out non-specific effects of dibucaine, we performed the following experiments. First, we repeated the dibucaine application in the presence of antagonists for voltage and ligand-gated channels that may be open near the -60 mV holding potential values (TTX 0.4 µM, TEA 1 mM, 4AP 1 mM, TTAP2 20 µM, NBQX 10 µM and gabazine 10 µM to target, respectively, voltage gated Na^+^, K^+^ and T-type Ca^++^ channels, and the AMPA and GABA_A_ ionotropic receptors). Under these conditions, a similar decrease in the HC was observed, from -18.1 ± 8.1 pA in control conditions to -10.8 ± 5.4 pA in the presence of the drug (**Figure 2Cc**, n = 11, p = 0.001), representing a 41 ± 12 % reduction. To note, there was no difference between the control HC recorded in the two conditions (without or with blockers, p = 0.9, Wilcoxon–Mann–Whitney test). Second, to rule out the possible contribution of conductances sensitive to the calcium flowing into the cell through PKD2L1 channels, we patched the neurons with the same intracellular solution + 10 mM BAPTA. Under these conditions, a decrease in the HC was still observed when pressure applying dibucaine, from -13.0 ± 5.7 pA in control conditions to -7.8 ± 2.5 pA ( **Figure 2Cd**, n = 11, p = 0.001), a 35 ± 15 % reduction. Again, there was no difference between the control HC measured in the 3 different conditions (without blockers vs BAPTA, p = 0.17; with blockers vs BAPTA, p = 0.19, Wilcoxon–Mann–Whitney test). Such a marked effect of dibucaine on the HC and the RMP can be explained by the fact that CSFcNs are electrotonic compact neurons with a high input resistance (IR). Indeed, the IR measured from 14 somatic recordings was 1.8 ± 0.46 GΩ (**Figure 2Ce**; consistent with previous published values in the turtle^21^, rat^3^ and mice^14,15^). As we will show later in this work, PKD2L1 channels are concentrated in the ApPr. Therefore, a better estimation of the impact of a PKD2L1-mediated current on the neuron’s membrane potential should be obtained by measuring the IR directly from the ApPr. To do that we made whole-ApPr recordings, where we measured an average IR of 2.3 ± 0.5 GΩ (n = 29), statistically higher than the somatic one (p = 0.002; **Figure 2Ce**). Finally, we took advantage of the fact that during the slicing procedure some ApPrs are separated from the rest of the cell; these will be referred to as “isolated” ApPr (iAPr). We succeeded to record from those iApPr, where we measured an IR that was even higher than the previous values (4.4 ± 1.2 GΩ, n = 8; **Figure 2Ce**; p = 6 × 10^-6^: iApPr vs soma; p = 4 × 10^-7^: iApPr vs whole-ApPr). Altogether, these results indicate that the spontaneous activity induced by PKD2L1 channels gives rise to both a phasic and a sustained current. Both phasic and sustained components of the PKD2L1-associated currents have marked effects on membrane potential as CSFcNs and, particularly, their ApPr, display very high IR values.

**Fig 2.**
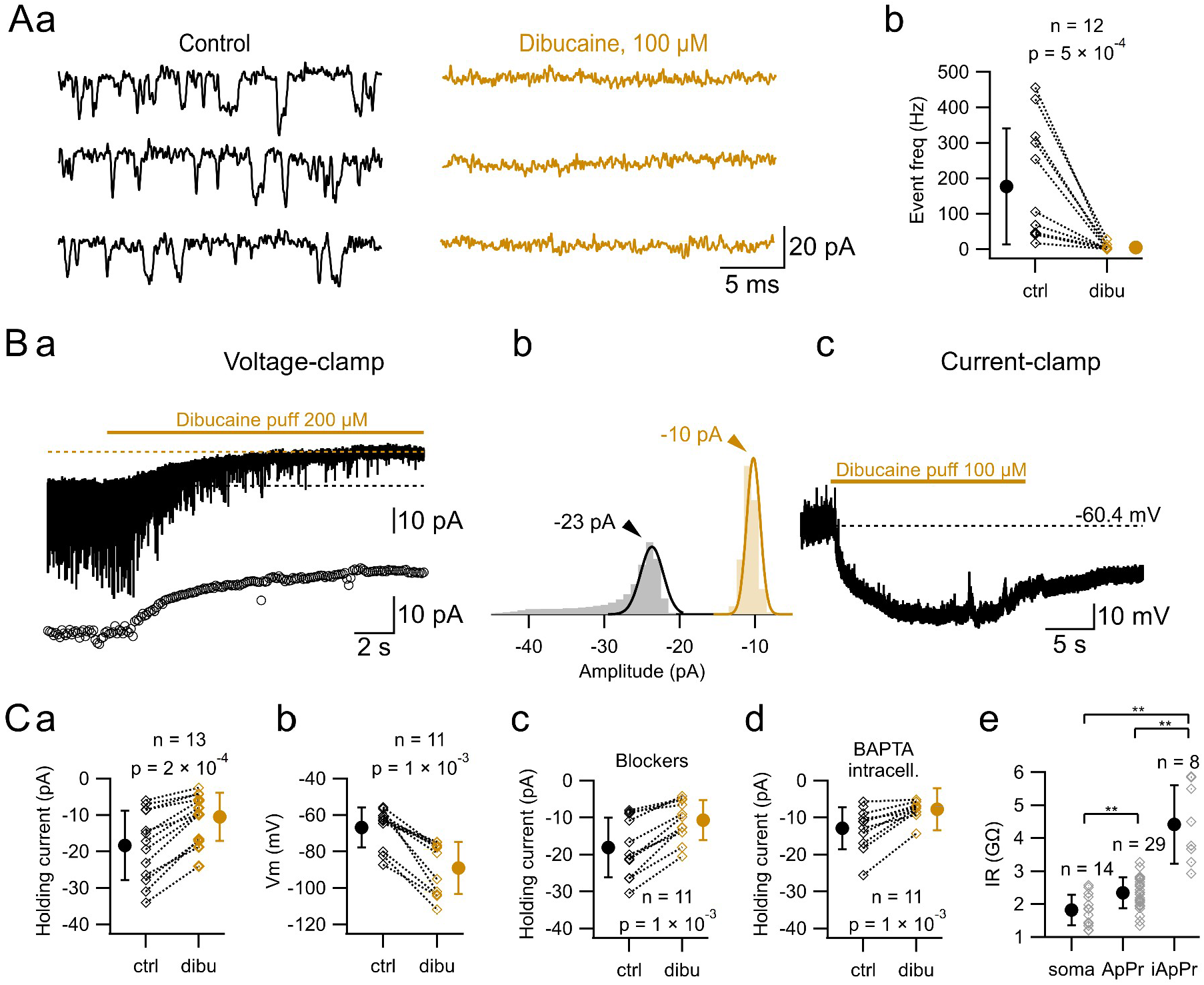
PKD2L1 channel activity mediates both phasic and tonic currents. **Aa.** Recording of a CSFcN in control condition and during pressure application of dibucaine hydrochloride (100 µM). **Ab.** Single channel event frequency calculated in control conditions and during dibucaine application. The average frequency was reduced from 176.9 ± 163 to 5 ± 8.3 Hz (n = 12, p = 5 × 10^-4^). **Ba.** Voltage-clamp recording showing how dibucaine application blocks the spontaneous events and reduces the holding current from -23 to -10 pA. The horizontal dotted lines indicate the average baseline current before and during dibucaine application. The bottom trace shows the average current value calculated from 100 ms time-periods, where the reduction in the holding current can be readily appreciated. **Bb.** Histograms of the recorded current during the control (black) and the dibucaine (orange) time periods. The data has been adjusted with a gaussian function (continuous lines). The mean values of the Gaussian fits correspond to the holding current values shown in **Ba** (dotted lines; -23 and -10 pA, control and dibucaine, respectively). **Bc.** Current-clamp recording showing the hyperpolarisation induced by dibucaine application, from -60.4 to -95 mV in this example. **a** and **c** correspond to different neurons. **Ca.** Effect of dibucaine pressure application on the HC: -18.3 ± 9.5 in control to -10.5 ± 6.6 pA in dibucaine (n = 13, = 2 × 10^-3^). **Cb.** Effect of dibucaine application on the resting membrane potential: -66.8 ± 11.0 in control to -89.0 ± 14.0 mV in dibucaine (n = 11, p = 1 × 10^-3^). **Cc**. Effect of dibucaine pressure application on the HC in the presence of voltage-gated and ionotropic channels blockers: -18.1 ± 8.1 pA in control to -10.8 ± 5.4 pA in dibucaine (n = 11, p = 1 × 10^-3^). **Cd.** Effect of dibucaine pressure application on the HC in the presence of 10 mM BAPTA in the internal solution of the recorded neurons: -13.0 ± 5.7 pA in control to -7.8 ± 2.5 pA in dibucaine (n = 11, p = 1 × 10^-3^). **Ce.** Input resistance (IR) values calculated from somatic (1.8 ± 0.46 Gfi, n = 14), ApPr (2.3 ± 0.48 Gfi, n = 29) and isolated ApPr (4.4 ± 0.12 Gfi, n = 8) recordings. p = 0.002, soma vs whole-ApPr; p = 6 × 10^-6^, iApPr vs soma and p = 4 × 10^-7^, iApPr vs whole-ApPr. In **Ab** and **C**, diamonds correspond to individual neurons and circles to the mean ± SD. Statistical comparison between groups was performed with a Wilcoxon signed-rank test for paired data (**Ab** and **Ca** to **d**) and a Wilcoxon–Mann–Whitney for unpaired data (**Ce**).

It has been reported that calmodulin modulates PKD2L1 channels through a direct interaction and that blocking calmodulin increases the activity of the channel^11,19,22^. In order to test whether this also applies for PKD2L1 channels in CSFcNs, we recorded channel activity under control conditions and in the presence of the calmodulin inhibitor calmidazolium. As expected, pressure-application of calmidazolium (10 or 20 µM) at the ApPr increased the channel activity, as shown in the representative recording in **Supplementary Figure 1Aa** (top panel). The increase in channel activity is shown in the middle panel, where the membrane charge is plotted as a function of time. It can be seen that the slope of the charge, which is dependent on the spontaneous openings of the channels, suddenly rises when calmidazolium is applied. The lower panel shows the normalized membrane charge (average *±* SD) of 10 different CSFcNs before, during and after calmidazolium application. In all neurons tested ( **Supplementary Figure 1Ab** for an example), calmidazolium application also increased the HC (from -13.3 ± 5.3 to -16.0 ± 4.7 pA, n = 10, p = 0.008, **Supplementary Figure 1Ac**), a 24 ± 20 % increase. This effect of calmidazolium on the HC had a dramatic effect on the RMP of the recorded cells. In I-clamp, calmidazolium induced a depolarization (**Supplementary Figure 1B**), from -62.3 ± 3.3 in control conditions to -50.0 ± 5.2 mV in the presence of calmidazolium, n = 10, p = 0.002, as well as an increase in the spontaneous EPSP frequency (lower part of **Supplementary Figure 1Ba**). This experiment shows that the increase in the activity of PKD2L1 channels impacts both phasic and sustained currents. This is consistent with our previous conclusion that blocking PKD2L1 channels likewise reduces both the phasic and the sustained component of PKD2L1-associated currents. Also, it highlights the importance of the intrinsic membrane properties, in this case the input resistance, in setting CSFcN excitability.

### Effect of extracellular pH on PKD2L1-mediated phasic and sustained currents

We next asked whether physiological stimuli can also modulate PKD2L1 activity in a similar fashion as dibucaine and calmidazolium. To address this issue we performed changes in extracellular pH, as CSFcNs have been shown to be sensitive to changes in extracellular pH (see Introduction and further below). When a pH 6.5, 10 mM HEPES-buffered extracellular solution was pressure-applied to the ApPr, a decrease in PKD2L1 activity was observed, as shown in **Figure 3Aa, top trace**, and in the corresponding inset (traces “7.4” and “6.5”). The decrease in channel activity can be appreciated in a plot of the current charge as a function of time (**Figure 3Aa, bottom trace**). **Figure 3Ab** shows the normalized current integral calculated from 10 different cells before, during and after the pressure application of the acidic solution. The dotted line represents a linear fit to the first 3 seconds of the recording, which corresponds to the control period and illustrates how the slope of the control period decreases during the application of the pH 6.5 solution. The decrease in channel phasic activity was also manifested as a decrease in the single channel open time during 500 ms (see the Materials and Methods section), from 46 ± 47 to 7 ± 7 ms (ctrl vs pH 6.5, n = 11, p= 0.001; **Figure 3Ca**) and a decrease in the n_max_ value, from 1.7 ± 0.60 to 1 ± 0 (ctrl vs pH 6.5, n = 11, p = 0.016; **Figure 3Cb**). Meanwhile, the single channel current amplitude was not affected (-14.7 ± 1.34 vs -14.0 ± 1.7, ctrl vs pH 6.5, n = 10, p = 0.15). Apart from the effects on the phasic currents, application of the acidic solution was also accompanied by a parallel decrease in the recorded HC, from -16.8 ± 6.7 to -12.0 ± 5.2 pA (**Figure 3Cc**, n = 11, p = 0.001), a 28.6 ± 12.7% reduction. The effect on the holding current can be seen in the VC recordings shown in **Figure 3Aa, top**, but is more clearly seen when the recorded current is averaged over 100 ms time periods (**Figure 3Aa, middle** graph). When the cell was held in I-clamp, pressure application of the pH 6.5 solution produced a marked hyperpolarization of the RMP (**Figure 3B** for an example), from -64.5 ± 5.2 to -76.0 ± 3.4 mV (n = 8, p = 0.008; **Figure 3Bd**). In 3 out the 8 experiments, a quickly desensitizing inward current was observed in VC at the onset of the application of the acidic solution (**Supplementary Figure 2Aa**), which produced a short-lasting depolarization in I-clamp that was followed by the persistent hyperpolarization described above (**Supplementary Figure 2Ab**). As previously described^14^, this current probably reflects the opening of ASIC channels. This current was not further characterized.

**Fig 3.**
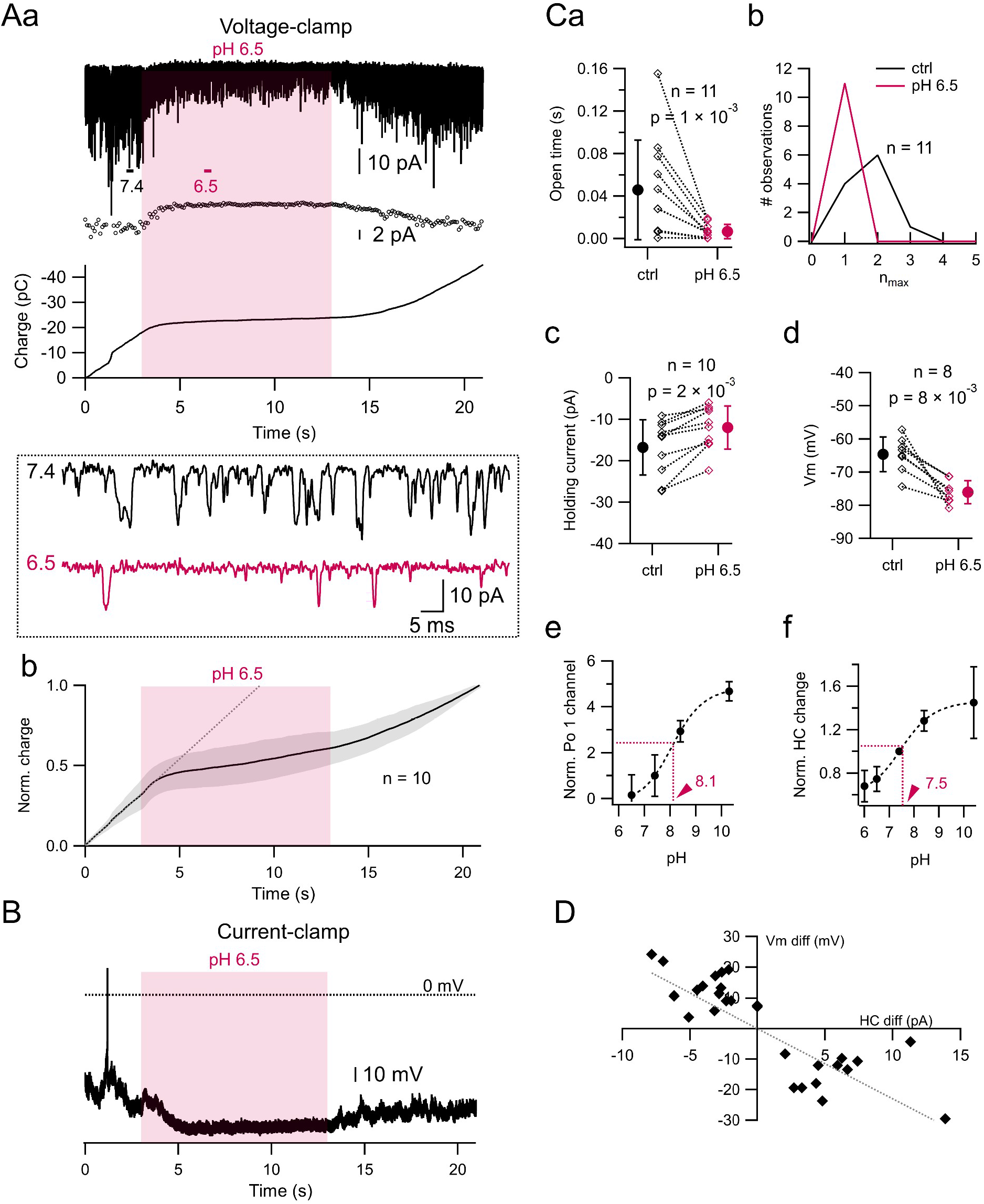
pH sensitivity of PKD2L1-mediated phasic and tonic currents. **Aa. Top.** Spontaneous PKD2L1 channel activity recorded in a CSFcN at a -60 mV holding potential. A pH 6.5 solution was pressure-applied during 10 s (starting at 3 s, red area). **Middle.** Holding current calculated from the recording shown on the top. **Bottom.** Membrane charge calculated from the recording shown on the top. **Inset.** Segments **7.4** and **6.5** in **Aa** are shown with expanded time and amplitude scales in order to see the decrease in the spontaneous single channel activity during the application of the acidic solution, without any change in the single channel current. **Ab.** Mean membrane charge ± SD calculated (gray surface) from 10 neurons tested in the same conditions as the cell shown in **a**. The dotted line corresponds to a linear fit to the control period (first 3 seconds of the recording). The application of the acidic solution produces a clear decrease in the slope of the membrane charge, indicating a decrease in the spontaneous openings of the channels. **A. B.** Same experiment as in **a**, but the spontaneous activity was recorded in I-clamp. The application of the acidic solution produces a hyperpolarization of the RMP (from -57.2 to -80 mV in this example). **Ca.** The single channel open time during 500 ms decreased from 46 ± 47 ms in control conditions to 7 ± 7 ms during the pressure application of a pH 6.5 solution (ctrl vs pH 6.5, n = 11, p= 0.001). **Cb.** Histograms of the n_max_ values calculated during 500 ms spontaneous, control recordings (black curve) and during the pressure application of a pH 6.5 solution (red curve). The mean n_max_ decreased from 1.7 ± 0.60 to 1 ± 0 (ctrl vs pH 6.5, n = 11, p = 0.016). A two sample Kolmogorov-Smirnov test also indicated a significant difference between the 2 distributions (p = 0.013). **Cc.** Effect of a pH 6.5 solution application on the holding current: -16.8 ± 6.7 in control conditions vs. -12.0 ± 5.2 pA in the pH 6.5 solution (n = 10, p = 0.001). **Cd.** Effect of a pH 6.5 solution application on the resting membrane potential: -64.5 ± 5.2 in control conditions vs. -76.0 ± 3.4 mV in the pH 6.5 solution (n = 8, p = 0.008). **Ce.** Normalized apparent po1 (apparent po1 test/apparent po1 at pH 7.4) as a function of pH. N = 10 for pH 6.5, 25 for pH 7.4, 8 for pH 8.4 and 7 for pH 10.4. The dotted line shows the fitting of the data to a Hill equation that yields a half pH value of 8.1 ± 0.04. Error bars are SDs. **Cf.** Normalized holding current (HC test/HC at pH 7.4) as a function of pH. N = 5 for pH 6, 5 for 6.5, 6 for pH 8.4 and 7 for pH 10.4. The dotted line shows the fitting of the data to a Hill equation that yelds a half pH value of 7.5 ± 0.02. Error bars are SDs. In **C**, diamonds correspond to individual neurons and circles to the mean ± SD. **A. D.** Relationship between Vm and HC differences recorded in 28 individual neurons. The dotted gray line is a constrained fit (passing through the origin) with a linear function (ax + b, where b = 0), which shows a slope of 2.3 ± 0.3 GΩ. The 2 variables are strongly correlated (Pearson linear correlation coefficient = -0.84; r^2^ = 0.71). In **C**, statistical comparison between groups was performed with a Wilcoxon signed-rank test.

On the other hand, when a pH 8.4 HEPES-buffered extracellular solution was pressure-applied, a clear increase in PKD2L1 activity was recorded (**Supplementary Figure 3A** and **B**). This was manifested as increases in the current integral (**Supplementary Figure 3A, bottom**), the mean single channel open time (from 16 ± 11 to 126 ± 73 ms; ctrl vs pH 8.4, n = 8, p = 0.078; **Supplementary Figure 3Da**) and the nmax value (from 1.25 ± 0.46 to 2.75 ± 0.70; ctrl vs pH 8.4, n = 8, p = 0.016; **Supplementary Figure 3Db**). As expected, there was also a slight increase in the HC (from -11.7 ± 7.3 to -15.2 ± 7.9 pA, n = 8, p = 0.008, **Supplementary Figure 3Dc**) and a depolarization of the RMP (from -75.2 ± 7.0 to -66.2 ± 10 mV, n = 6, p = 0.03, **Supplementary Figure 3C** and **Dd**). The single channel current remained unchanged (-16.85 ± 0.95 vs -17.3 ± 1.0 pA, ctrl vs pH 8.4; respectively, n = 8, p = 0.45).

The experiments presented so far indicate that the basal activity of PKD2L1 channels plays a central role in regulating CSFcNs excitability. This regulation takes place at two levels: by a modulation of the phasic activity of the channels, as has already been described^14,15^, and also by a modulation of a novel PKD2L1-dependent sustained current that has a dramatic effect in setting the RMP in CSFcN. In this sense, PKD2L1 basal activity determines an excitability set-point that can be up and down-regulated by small pH changes. This is clearly seen when plotting the apparent po (normalized relative to the apparent po at pH 7.4) as a function of pH (**Figure 3Ce**). The experimental data has been fitted by a Hill equation that shows a pH half-value of 8.1 ± 0.04, very close to physiological pH and to previously estimated values in expression systems^23^ and computational model^24^. A similar result is obtained when plotting the holding current change (in relation to the control value at pH 7.4) as a function of pH. Fitting the data with a Hill equation gives a pH half-value of 7.5 ± 0.02 (**Figure 2Cf**). In order to better appreciate the effect of the HC on the Vm, we calculated the Vm and the HC differences produced (in individual neurons) by the different experimental challenges tested here, and then plotted the Vm vs the HC difference. The plot, shown in **Figure 3D**, indicates a strong correlation between the 2 values. The fit of the data with a linear function (dotted gray line) indicates a slope value of 2.3 GΩ, very close to the IR values calculated for the ApPr (**Figure 1Ee**). Taken together, these data indicate that both the PKD2L1 channel po and the holding current are very sensitive to extracellular pH changes, that the maximal pH sensitivity occurs within the physiological range and that small changes in the sustained current can have a substantial effect on CSFcNs RMP (≈ 2.3 mV/pA).

### The use of laser photolysis to assess the pH sensitivity of CSFcNs

It has been reported that the sensitivity of CSFcNs to pH changes depends on the activity of two different channels, PKD2L1 and ASICs. Those studies used a combination of electrophysiological methods with bath or pressure applications of drugs, but these strategies suffer from limited spatial resolution, preventing precise localization of channel activity within the CSFcN membrane. To overcome this limitation and to precisely map the sub-cellular sensitivity to pH of the different CSFcN compartments, we employed laser photolysis, as this approach provides high spatial resolution (the laser can be focused to a small spot, with x-y dimensions of 1.5 µm^25,26^) as well as a high temporal resolution (the laser can be switched on and off very rapidly, thus producing almost instantaneous agonist changes). This is shown schematically in **Figure 4A-B**. Briefly, after the different CSFcN compartments (soma, dendrite and ApPr) were identified by fluorescence of Alexa 594, the focus of the objective was positioned in the targeted compartment (ApPr in the example in **Figure 4A**), where laser photolysis was evoked.

**Fig 4.**
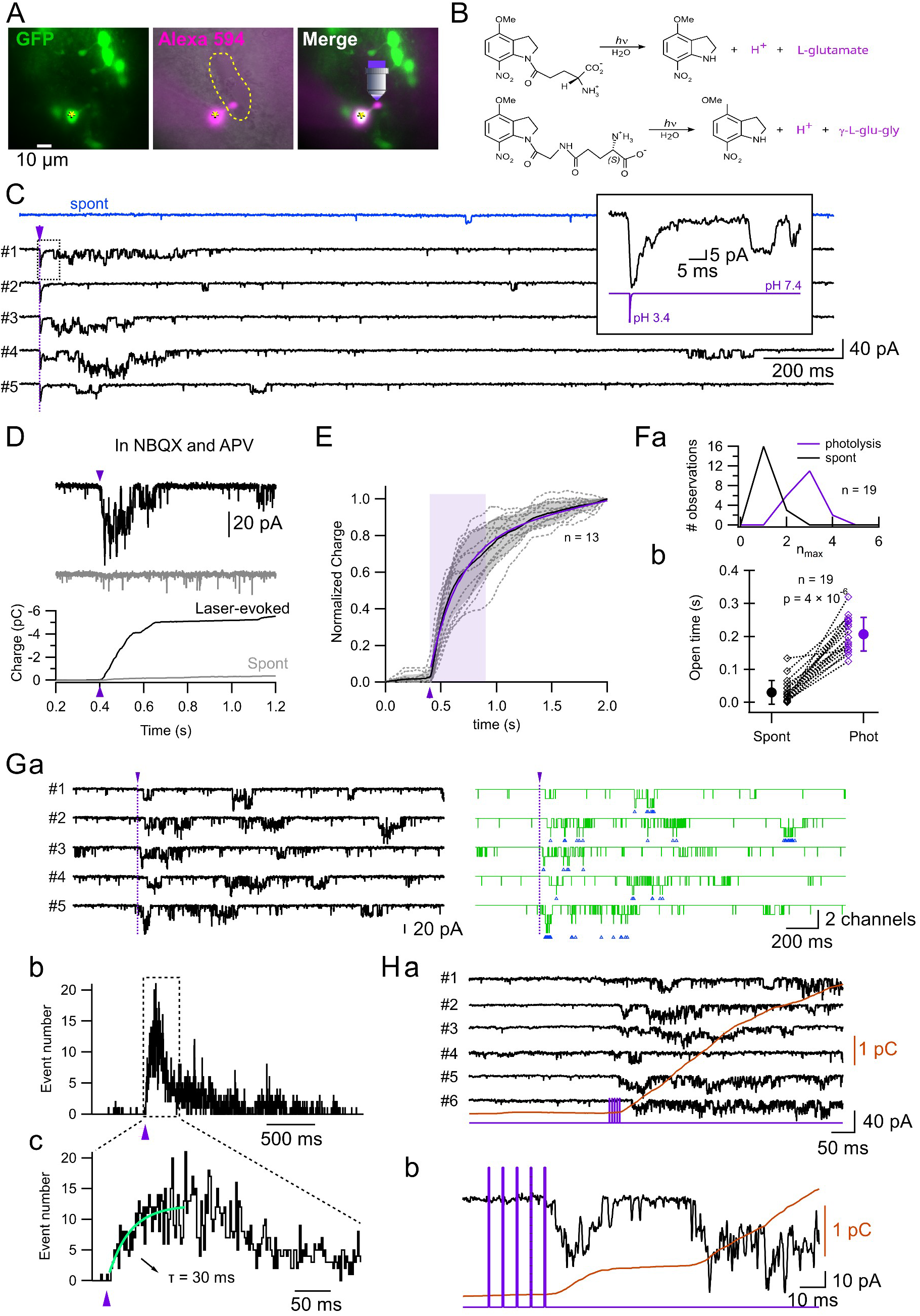
Proton photolysis induces a recovery current in CSFcNs. **A.** Schematic of the experiment. Pictures showing the eGFP fluorescence (left), the Alexa 594 fluorescence (middle) and the merging of the 2 channels (right). The yellow star indicates the recorded cell, and the objective the location of the targeted compartment. The yellow dotted line shows the approximate boundaries of the central canal. **B.** Schematic of the photolysis reaction shown for the 2 caged compounds used: MNI-Glutamate (top) and MNI-γLGG (bottom). **C.** 2 second-long recordings showing the typical response of a CSFcN (shown in **A**) to the photolysis of MNI-Glutamate on the ApPr (black traces). The photolysis was repeated 5 times with 10 seconds intervals. The blue, upper trace, shows the spontaneous recording (no photolysis) for comparison. The inset shows sweep #1 in an expanded scale in order to appreciate the fast kinetics of the AMPA_R_-mediated current. Vertical, magenta arrowhead and dotted line indicate the laser pulse (1 ms, 4.3 mW; which is measured with a photodiode in the laser path). **D. Top.** The black trace shows the current evoked by the photolysis of MNI-γLGG with a 0.5 ms, 4.3 mW laser pulse on the ApPr, and the gray trace the spontaneous current recording in the same CSFcN. During the spontaneous recording the maximum number of channels opened simultaneously (n_max_) was 1, and during the 500 ms window after the photolysis n_max_ was 3. The total open time in a time window of 500 ms was 17 ms during the spontaneous recording and 196 ms after the photolysis. **Bottom.** Membrane charge calculated from the above recordings. The vertical, magenta arrowhead indicates the laser pulse. **E.** Normalized (to the 2 s value) membrane charge as a function of time. Black trace shows the average, gray area the ± SD and dotted traces individual experiments (n = 13). The magenta continuous line represents the fit of the average curve with the sum of a linear + an exponential function representing the increase evoked by the photolysis and the linear increase due to the spontaneous channel openings, respectively (see methods). The τ of the exponential function was 258 ± 2 ms. The x-span of the magenta area represents 500 ms (≈ 2 τs). The vertical, magenta arrowhead indicates the laser pulse. **Fa.** Histograms of the n_max_ values calculated during 500 ms time windows for spontaneous recordings (black curve) and after photolysis (magenta curve). The mean n_max_ was 1.16 ± 0.37 in spontaneous recordings and 2.8 ± 0.63 after photolysis (n = 19, p = 4 × 10^-6^). A two sample Kolmogorov-Smirnov test also indicated a significant difference between the 2 distributions (p = 1 × 10^-6^). **Fb.** Total open time for a single channel during a 500 ms time window (spontaneous vs photolysis). The mean single channel open time increased from 30 ± 36 to 207 ± 51 ms (500 ms spontaneous recording vs 500 ms after the photolysis pulse; n = 19; p = 4 × 10^-6^). **Ga.** Example of laser-evoked currents (left, black traces) and the corresponding idealized events (right, green traces). MNI-γLGG was used in this experiment. The blue arrowheads below each idealized trace indicates the timing of double events (where 2 channels opened simultaneously). Vertical, magenta arrowheads and dotted lines indicate the laser pulse (1 ms, 4.3 mW). **Gb.** Latency distribution of double events (95 photolysis repetitions from 20 neurons) shown with 2 different time resolutions (the graph on the right corresponds to the time indicated by the dotted rectangle on the graph on the left). The rising phase of the histogram has been fitted with an exponential function (green trace) that has a τ of 30 ± 12 ms. The magenta arrowheads indicate the timing of the laser pulse. **Ha.** The black traces show the responses of a CSFcN to the photolysis of MNI-γLGG on the ApPr with a train of 5 laser pulses (2 mW, 0.5 ms duration) at 180 Hz. The photolysis train was repeated 6 times (#1 to #6) with 10 seconds intervals. The brown curve shows the membrane charge calculated from the average of the 6 sweeps. **A. b.** Sweep #2 and its corresponding membrane charge are shown on an expanded time scale in order to better appreciate the onset of the response, that happens after the end of the train. In **F**, statistical comparison between groups was performed with a Wilcoxon signed-rank test.

To induce pH changes we used the glutamate cage MNI-glutamate, as its photolysis generates the stoichiometric release of one glutamate molecule together with one proton^27^ (**Figure 4B, top** reaction). MNI-glutamate uncaging in the ApPr of CSFcNs elicited two currents with different characteristics. A first component had very quick rise and decay times (values of the example shown in **Figure 4C**: amplitude 38 pA, 10-90 % risetime 0.86 ms and decay time constant 2.6 ms; see inset for details). It appeared immediately after the laser pulse (magenta arrowhead in **Figure 4C**) and fluctuated little among repetitions. A second component was indistinguishable from the PKD2L1-dependent spontaneous activity recorded above. It followed the laser pulse with variable latencies (although always longer than the latencies of the first component) and fluctuated widely among repetitions. The first current was blocked by AMPAR antagonists (NBQX or CNQX) and was therefore attributed to AMPA receptor activation. The second component of the response was spared by both AMPA or NMDAR blockers (the experiment shown in **Figure 4D** was done in the presence of NBQX and APV) and was presumed to reflect the pH change. To confirm that this current resulted from the release of protons (and not glutamate), we used the caged compound MNI-γLGG, which has the same photochemistry as MNI-glutamate except that it releases the inactive enantiomer of the AMPA_R_ antagonist γDGG^28^ together with a proton (**Figure 4B, bottom** reaction). This failed to produce the short latency current, but produced the same long latency current as above, indicating that the late current is related to proton release rather than glutamate ( **Figure 4G** and **H** for representative examples). Also, no current could be observed when the ApPr was illuminated in the absence of a caged compound (data not shown), indicating that the current is not due to a light artifact or to photodamage.

### The photolysis-induced current is mediated by PKD2L1 channels

Inspection of the experimental traces presented in **Figure 4C** suggests that the photolysis of an MNI-cage in the ApPr of CSFcNs increases the activity of PKD2L1 channels with a longer latency than that of AMPA-Rs mediated currents. The increase in channel activity was analysed in different ways. First, the membrane charge was calculated from spontaneous recordings (gray traces in **Figure 4D**) and from recordings where MNI-glutamate (or MNI-γLGG) was photolysed (black traces in **Figure 4D**). As already mentioned (**Figure 3**), the membrane charge calculated from spontaneous sweeps shows a linear increase as a function of time, whereas that calculated from laser-evoked sweeps shows a sudden slope increase induced by the laser pulse, that recovers later

(**Figure 4D, bottom plot**). **Figure 4E** shows the average normalized membrane charge (black trace) calculated from 13 different neurons where photolysis was evoked on the ApPr. The data was fitted with the sum of a linear and exponential functions. The time constant of the exponential function, τ, is equal to 258 ms. The magenta-shaded area in the graph (in **Figure 4E**) spans 500 ms (≈ 2 τ_s_), representing the estimated lifetime of the photolysis-evoked increase in PKD2L1 channel activity.

Second, we measured the maximum n (n_max_; maximum number of open channels) during spontaneous recordings and during the 500 ms period after the photolysis pulse. This analysis indicated that n_max_ systematically increased from 1.16 ± 0.4 to 2.8 ± 0.6 (500 ms spontaneous recording vs 500 ms after the photolysis pulse; n = 19; p = 4 × 10^-6^). In the example shown in **Figure 4D**, n_max_ increases from 1 to 3. **Figure 4Fa** shows the distribution of n_max_ in the two conditions.

Finally, we measured the total open time of a single channel during the control period (500 ms of spontaneous recording) and during the 500 ms time window after the photolysis pulse. This analysis revealed a dramatic increase in the channel open time, from 30 ± 36 to 207 ± 51 ms (500 ms spontaneous recording vs 500 ms after the photolysis pulse; n = 19; p = 4 × 10^-6^; **Figure 4Fb**). In the example shown in **Figure 4D**, the open time increased from 17 to 196 ms. Altogether, the evidence presented in **Figure 4D** to **F** indicate that the apparent single channel open probability increases as a result of the photolysis.

As previously mentioned, the increase in channel activity does not happen during the laser pulse but a few milliseconds afterwards. In this sense, the latency difference between the AMPA-dependent and proton dependent components is striking. This difference can be clearly appreciated in the inset in **Figure 4C**, where the AMPA-dependent component coincides with the laser pulse (magenta trace) whereas the proton-dependent component appears more than 30 ms later. The PKD2L1 current latency in sweep #4, which is the shortest among the 5 repetitions, is 11 ms. To characterize the latency distribution of the PKD2L1-mediated single currents relative to laser pulse timing, we measured the current onset times from idealized traces during 2-second recording periods (see **Figure 4Ga** for an example; experiments done in continuous presence of NBQX or with MNI-γLGG). To minimize contamination from spontaneous channel openings, we only considered double events (simultaneous opening of two channels) as the spontaneous occurrence of such events is very low. From the onset times of these double events, we constructed the latency distribution histogram shown in **Figure 4Gb**. The probability of 2 channels opening at the same time gradually rises after the laser pulse with a time constant of 30 ± 12 ms (**Figure 4Gc**), peaking at 108 ms after the laser pulse (while the AMPA current peaks a few µs after the laser pulse; see inset in **Figure 4C**). This analysis, together with our calculation that photolysis induces a transient pH drop of around 4 units (from 7.4 to ≈ 3.3) that returns to baseline extremely fast (see materials section), suggests that the photolysis-evoked PKD2L1 current represents a pH dependent recovery current (that follows the cessation of acidification). Indeed, it has been shown that PKD2L1 channels, which are activated by alkali and inhibited by acid, are also activated by acid removal ^24,29,30^, giving rise to a recovery response.

In order to confirm this, we designed and experiment where instead of applying a single laser pulse, we applied a train to acidify the local environment. If the photolysis-evoked current is indeed a current that follows the cessation of acidification, then the majority of the single-channel events should appear at the end or after the train. This is indeed what happens, as shown in **Figure 4Ha**. In this cell, a photolysis train (5, 500 µs pulses at 180 Hz) was applied 6 times on the ApPr, and the majority of the single channels events appear after the end of the pulse. This can be more clearly seen from the charge (calculated from the average current) and on sweep #2 (**Figure 4Hb**) shown on an expanded time-scale.

The experiments presented so far strongly suggest that the photolysis-evoked current is mediated by PKD2L1 channels. To confirm this, we performed the following experiments/analysis. Firstly, channel activity was blocked when the photolysis is performed in the continuous presence of dibucaine. This can be appreciated in the representative trace shown in **Figure 5A**: in the presence of dibucaine (gray trace), both the spontaneous currents and the laser-evoked currents were blocked. The bottom panel shows the time-dependent current integrals calculated from the upper traces. **Figure 5B** shows the average normalized membrane charge recorded under two conditions, control (n = 13 neurons, same graph as in **Figure 4E**) and dibucaine (n = 6 neurons). Some of the experiments in the presence of dibucaine were done in the absence of AMPAR blockers. In these cases, a clear fast, inward, AMPAR-mediated current can be appreciated which is not followed by any single channel event (**Figure 5D**), indicating that the lack of response was not due to an unresponsive or damaged ApPr.

**Fig 5.**
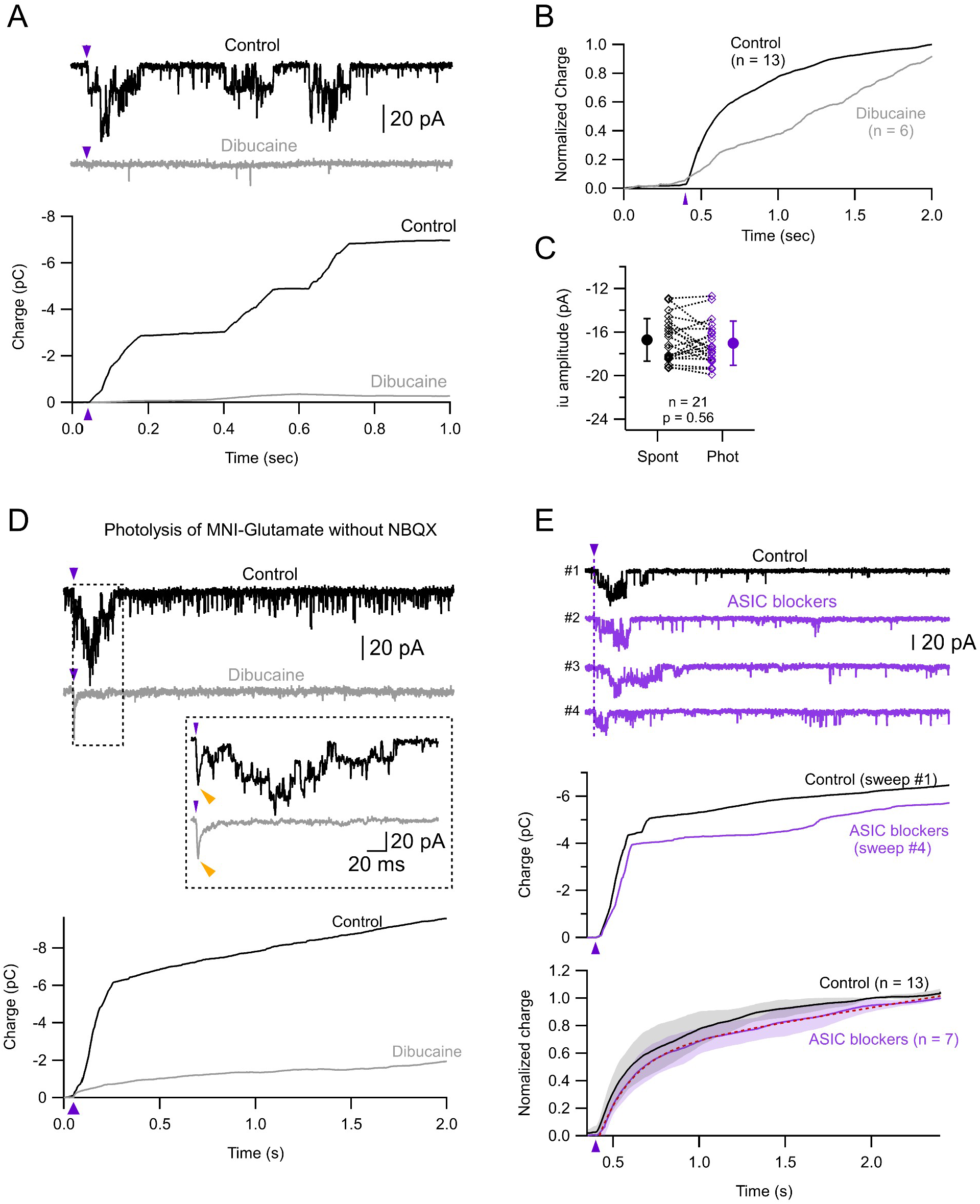
Currents induced by proton photolysis in the ApPr depend on PKD2L1 and not ASIC channels. **A. Top.** Example of laser-evoked currents in control conditions (black trace) and in the presence of 100 µm dibucaine (gray trace). **Bottom.** Membrane charge calculated from the recordings shown above. The vertical, magenta arrowhead indicates the laser pulse (1 ms, 4.3 mW). **B.** Average, normalized membrane charge calculated from recordings in control conditions (same data as in **4E**) and in the presence of dibucaine. The vertical, magenta arrowhead indicates the laser pulse. **C.** Single channel current calculated from spontaneous current recordings (spont) and during a 500 ms time window after the photolysis. Gray symbols correspond to individual neurons. Mean ± SD (black symbols): -16.7 ± 2.0 (spont) to 17.0 ± 2.0 (phot) pA, n = 21 neurons, p = 0.56. Wilcoxon signed-rank test. **D. Top.** Example of laser-evoked currents induced by the photolysis of MNI-Glutamate without NBQX (black trace) and without NBQX + dibucaine (gray trace). The inset shows the current traces in expanded scales to better illustrate the AMPA-mediated current (yellow arrowheads). **Bottom.** Membrane charge calculated from the above recordings. The vertical, magenta arrowhead indicates the laser pulse (1 ms, 5.2 mW). **E. Top.** Example of laser-evoked currents induced by the photolysis of MNI-Glutamate without (black trace) and with ASIC channel blockers (magenta traces). **Middle.** Membrane charge calculated from sweeps number 1 (control) and 4 (ASIC blockers). **Bottom.** Normalized (to the 2 s value) membrane charge as a function of time in control conditions (black trace, n = 13; same data as in **Figure 4E**) and in the presence of ASIC blockers (magenta trace, n = 7). The dotted red line represents the fit to the data with the same function as in **Figure 4E**. The τ value in the presence of ASIC blockers was 190 ms, close to the 258 ms value for the control curve. Shaded areas represent ± SDs. The vertical, magenta arrowhead indicates the laser pulse (500 µs, 5.8 mW). In **C**, statistical comparison between groups was performed with a Wilcoxon signed-rank test.

Secondly, the single channel current amplitude evoked by the laser pulse (calculated during the 2 τ_s_ time window after the laser pulse; see above) was indistinguishable from that obtained during spontaneous current activity recordings (**Figure 5C**, -16.7 ± 2.0 to 17.0 ± 2.0 pA, n = 21, p = 0.56).

ASICs have been shown to be present in CSFcNs^13–15,31^. To rule out any contribution of ASIC to the photolysis-evoked response on the ApPr, we performed the experiment in the continuous presence of the ASIC blockers PsTx (100 nM) and ApeTx2 (100 nM). In these conditions, neither the single channel events nor the charge increase evoked by photolysis were affected (**Figure 5E**), confirming that the increase in channel activity related to the release of protons by photolysis on the ApPr is dependent on the activation of PKD2L1 channels.

### The PKD2L1-mediated current is generated in the ApPr

The next series of experiments was designed to assess the spatial sensitivity of CSFcNs to pH changes. To this aim, we photolysed MNI-compounds in different CSFcNs compartments. **Figure 6A** shows the result of an experiment where the photolysis of MNI-γLGG was performed either in the ApPr, the soma or 5 µm away from the ApPr (**Figure 6Aa**). As expected, photolysis in the ApPr elicited the photolysis-evoked current described in the previous section, characterized by the increase in the unitary activity of PKD2L1 channels (**Figure 6Ab**, black traces) and the corresponding increase in membrane charge; on the contrary, photolysis in the soma ( **Figure 6Ab**, pink traces) or in the dendrite (data not shown) did not produce a sizable increase in the unitary currents. **Figure 6Ac** summarizes the photolysis-evoked changes on the membrane charge. Photolysis on the ApPr elicited a pronounced increase in membrane charge, whereas photolysis in the soma produced no detectable change. Notably, the response to photolysis in the soma was indistinguishable from the photolysis 5 µm away from the ApPr (magenta vs green traces, respectively), which is more evident in the bottom panel showing the same data set in an expanded amplitude scale.

**Fig 6.**
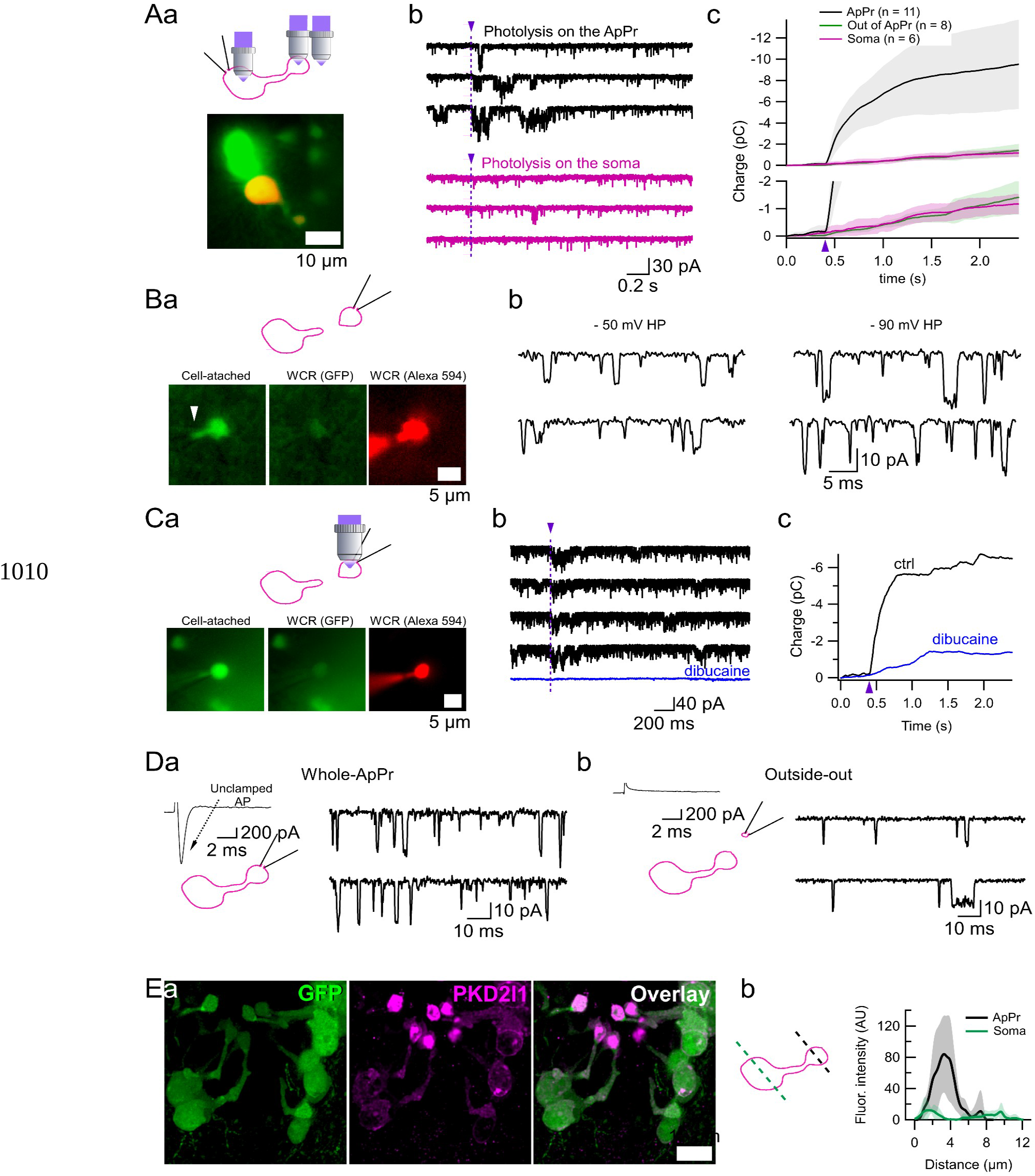
PKD2L1 channels are segregated to the ApPr. **Aa. Top. Schematic** drawing showing the different photolysis locations. **Bottom.** Superimposition of pictures showing the eGFP (green) and the Alexa 594 (red, which corresponds to the recorded cell fluorescence. **Ab.** Representative sweeps showing the membrane currents recorded when the photolysis of MNI-γLGG was performed either on the ApPr (black traces) or on the soma (magenta traces; laser pulse duration = 700 µs; power 5.3 mW). Scale bars are the same for black and magenta traces. **Ac.** Average membrane charge calculated from recordings corresponding to the photolysis of MNI-γLGG or MNI-glutamate in different locations. Shaded areas correspond to ± SDs. Bottom graph shows the same traces but in a different amplitude scale. The fact that no PKD2L1-mediated activity can be triggered when the photolysis is done a few microns away from the ApPr (5 to 7 µm) is compatible with the high spatial resolution of the photolysis technique and the restricted proton diffusion. **Ba. Top.** Schematic drawing of experimental design. Whole-cell recordings were performed from iApPrs. **Bottom.** Pictures showing the different recording configurations: **left** in cell-attached, where the eGFP green signal can be seen inside the recording pipette as the ApPr membrane goes into the pipette when the positive pressure used for patching is released; **midddle** in whole-cell a few seconds after break-in, where it can be seen that the green eGFP fluorescence has already washed-out; **right** in whole-cell, where the morphology of the isolated ApPr can be appreciated by the red Alexa 594 fluorescence. **Bb.** Whole-cell recordings from the isolated ApPr shown in **a** at two different holding potentials, -50 and -90 mV. A clear PKD2L1-dependent spontaneous activity can be seen. HP: holding potential. Scale bars are the same for the currents recorded at the two holding potentials. **Ca. Top.** Schematic drawing of experimental design. The photolysis was produced on an isolated ApPr. **Bottom.** Pictures showing the different recording configurations. Same as **Ba**. **Cb.** Whole-cell recordings from the isolated ApPr shown in **a** and its response to photolysis. The black traces are in control conditions and the bottom, blue trace in the presence of 100 µM dibucaine (laser pulse duration = 1 ms; power 5.1 mW). **Cc.** Average membrane charge calculated from 4 individual sweeps in control and 4 individual sweeps in dibucaine. **A. D.** Recordings from an ApPr in the whole-cell configuration (**a**) and in the outside-out configuration (**b**). The traces on the left show the current responses to a 50 ms depolarization to -10 mV from a -60 mV holding potential. This voltage pulse induces an unclamped sodium spike in whole-cell but no response in outside-out, confirming that the outside-out patch has completely detached from the ApPr (as isolated ApPr do not have sodium currents). PKD2L1 activity is still present in the outside-out recording. **Ea.** Maximum intensity projection of 41 deconvolved optical sections spaced by 130 nm. CSFcNs expressing eGFP (green, left panel), which also express PKD2L1 receptors (magenta, middle panel.). The overlay of the 2 channels is shown on the right panel. The dotted white line in the left, GFP panel, indicates the distance along which the density plots shown in **b** were constructed. **Eb.** Density plots depicting anti-PKD2L1 mean fluorescence intensity (trace) ± SDs (shade) measured at the soma (green) and in the ApPr (black). n = 6 for each compartment. The fluorescence mas measured along the dotted green and black lines shown in the scheme. The maximal mean intensity is 84 for the ApPr and 12.5 for the soma. Brightness and contrast were adjusted for display purposes. Unsaturated images were used for quantification. In **A** and **C**, the vertical dotted lines and magenta arrowheads correspond to the timing of the photolysis pulse.

To further assess the sensitivity of ApPrs to pH changes, we exploited the fact that during the slicing procedure some ApPrs are separated from the rest of the cell. We term these cell fragments iApPrs. The iApPr were filled with Alexa 594 in order to confirm that they were indeed detached from the rest of the neuron (**Figure 6Ba**). Interestingly, iApPrs show single-channel currents with the same single-channel conductance and voltage-dependence as non-isolated ApPrs, as can be appreciated in the representative example shown in **Figure 6Bb**. Furthermore, iApPrs respond to proton photolysis similarly to non-isolated ApPrs (**Figure 6C**) and both the spontaneous single-channel activity as well as the photolysis-induced charge increase are blocked by dibucaine (**Figure 6Cb-c**).

To confirm the presence of PKD2L1 channels in ApPrs, we performed outside-out recordings from patches of membrane taken from the ApPr (**Figure 6D**). We first recorded the ApPr in the whole-ApPr configuration and then slowly took out the pipette until no unclamped action current could be seen (as iApPrs do not show voltage-gated sodium currents; **Figure 6Da-b**). PKD2L1 channel openings were observed in the outside-out configuration (**Figure 6Db**), albeit with a lower opening frequency than in the whole-cell configuration (**Figure 6Da**). Such single-channel activity was observed in 6/10 outside-out recordings.

Finally, we performed immunohistochemical experiments to characterise the location of the PKD2L1 protein in Gata3^eGFP^. As seen in **Figure 6E**, the PKD2L1-related fluorescence intensity (shown in magenta in **Figure 6Ea**) is >6-fold higher in the ApPr than in the soma (**Figure 6Eb**) or dendrite (not shown). The mean fluorescence in the ApPr was 84 *±* 49 AU and dropped to 12.5 *±* 8 AU in the soma. Altogether, our experiments indicate that the PKD2L1 channels are specifically enriched in the ApPr. Although we cannot rule out the presence of PKD2L1 channels in other compartments, our electrophysiological and immunohistochemical experiments suggest that functional PKD2L1 channels are topologically segregated to the ApPr.

### The ApPr and soma are strongly coupled

We have shown so far that PKD2L1 channels are predominantly located in the ApPr. Also, the possibility of recording from iApPrs led us to a serendipitous finding: ApPrs do not have active sodium conductances. As it was shown before, a single PKD2L1 channel opening can led to spiking in CSFcNs^15^. Taken together, this suggests that the voltage changes that occur in the ApPr need to propagate very efficiently down the soma (and probably the axon) in order to have an impact on the spike triggering zone.

In order to measure directly the degree of coupling, we performed paired recordings between the ApPr and the soma (for specific details on the paired recordings see the Materials and Methods section). Both compartments were recorded in I-clamp in order to measure the steady-state coupling coefficient (DC CC; see materials and methods section). In these conditions (**Figure 7A**), the DC CC when injecting current in the soma (**Figure 7Aa** for an exemplary experiment) was 0.9 ± 0.09 (n = 6 pairs; **Figure 7Ad**). The DC CC in the other direction was 0.92 ± 0.09 (the same 6 pairs; **Figure 7Ab** and **d**). We also recorded the spontaneous activity in both compartments, as the majority of the spontaneous potentials recorded in the soma reflect the unitary PKD2L1 single channel activity. The spontaneous events CC (ssp CC; see Materials and Methods section) recorded from the 6 pairs was 0.88 ± 0.05, similar to the DC CC measured with rectangular current injections. These experiments indicate that the spontaneous events originating in the ApPr travel to the somatic compartment with almost no attenuation.

**Figure 7.**
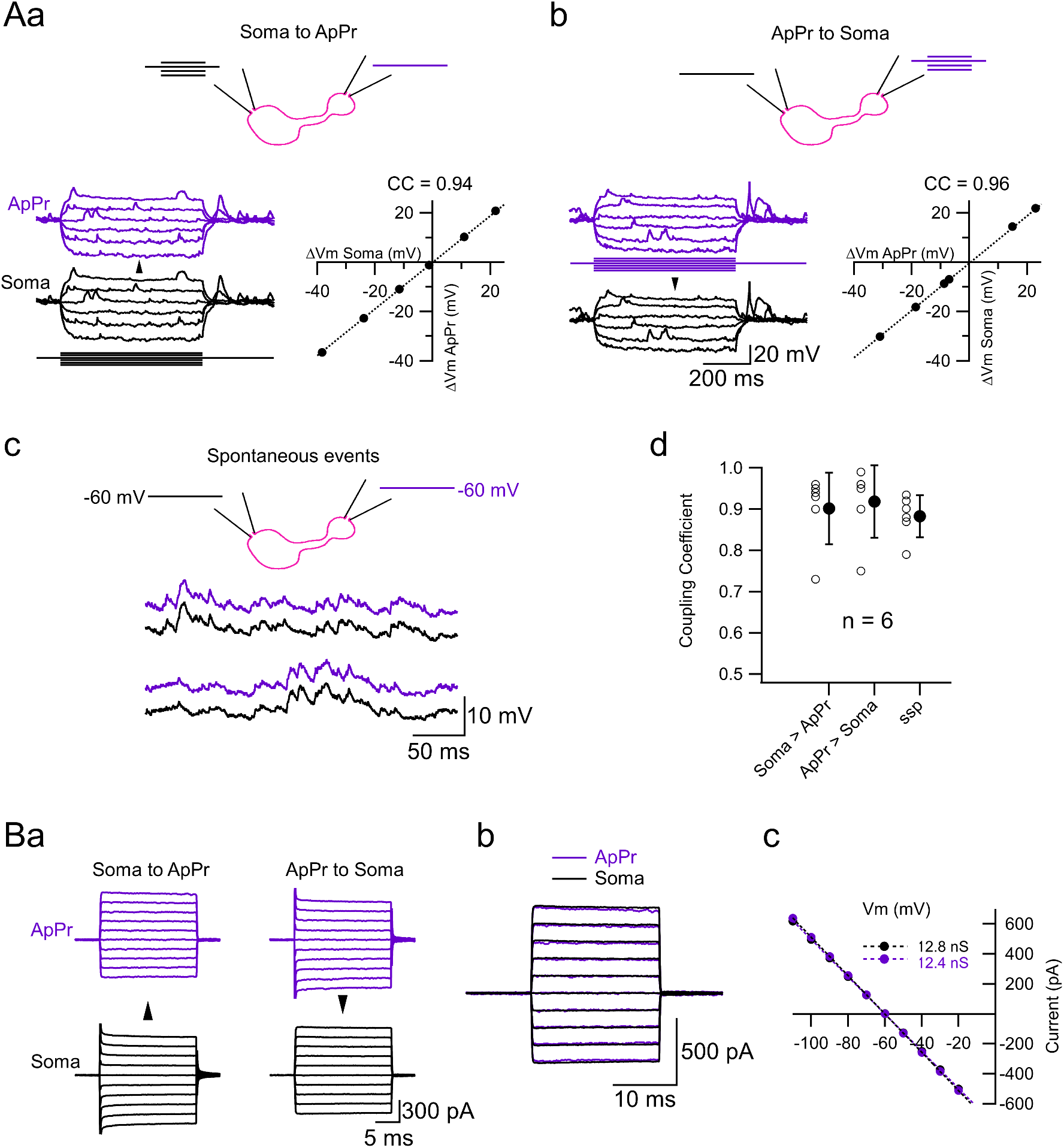
The coupling between the ApPr and the soma measured with paired recordings. **A.** Paired current-clamp recordings between the ApPr and the soma. The cartoons on top of the traces show the recording configurations. **Aa.** The current pulses are applied to the soma, and voltage measured both in the soma and the ApPr (**left**). The CC (0.94) is calculated by plotting the membrane voltage responses in the ApPr vs the membrane voltage responses in the soma (**right**). **Ab.** Same as in **a**, but now the current pulses are applied to the ApPr. The CC is 0.96. **Ac.** Spontaneous events recorded simultaneously in the ApPr and the soma. Resting membrane potential was -60 mV in both compartments. **Ad.** Coupling coefficient measured in 3 conditions: somatic current injection (0.9 ± 0.09), ApPr current injection (0.92 ± 0.09) and for spontaneous activity (soma/ApPr: 0.88 ± 0.05). **B.** Paired voltage-clamp recordings between the ApPr and the soma. **Ba.** Voltage pulses (-110 to -20 mV in 10 mV steps) are applied either to the soma ( **left**) or to the ApPr (**right**). **Bb.** Currents recovered in the non-pulsed compartments. **Bc.** The currents shown in **b** as a function of the voltage steps. The slope of each curve represents the coupling conductance between the 2 compartments (12.85 and 12.4 nS in this example). The average coupling conductance was 13.7 ± 0.62 (pulses applied to the ApPr) and 13.6 ± 0.62 nS (n = 7). All the recordings shown in this figure correspond to the same CSFcN.

Finally, we also recorded the 2 compartments in voltage-clamp in order to directly measure the coupling conductance between the ApPr and the soma. Both compartments were held at -60 mV, and a family of voltage pulses was applied to either of them (**Figure 7Ba**). In these conditions, the current recorded from the non-pulsed compartment should be identical in both directions (**Figure 7Bb**) and, most importantly, gives a direct measure of the conductance between the ApPr and soma^32^. In the example shown in **Figure 7B**, the coupling conductance measured when pulsing the ApPr (left column in **Figure Ba**) was 12.8 nS (**Figure 7Bc**) and that measured when pulsing the soma (right column in **Figure Ba**) was 12.4 nS (**Figure 7Bc**). The mean coupling conductance measured from 7 ApPr-soma pairs was 13.7 ± 0.62 (pulses applied to the ApPr) and 13.6 ± 0.62 nS (pulses applied to the soma), with a mean conductance ratio of 0.99 ± 0.03.

## Discussion

The pH sensitivity of neurons depends on both their anatomical features and the functional properties of their ion channels. As all neurons exhibit some degree of pH sensitivity^33^, investigating it needs the use of targeted techniques that can accurately assess both the anatomical and physiological properties of the channels involved in the response to pH. Our data, derived from the integration of whole-ApPr and outside-out recordings, along with laser photolysis of protons and immunohistochemistry, demonstrate that the unique ability of CSFcNs to function as precise CSF pH sensors arises from the distinct physiological properties of PKD2L1 channels and, crucially, their spatial segregation within the ApPr. These finding highlight how specialized anatomical and physiological features enable selective pH sensing.

From the anatomical standpoint, the functional evidence gathered here by combining whole-ApPr and outside-out recordings with laser-photolysis of protons and immunohistochemistry indicates that PKD2L1 channels are predominantly located in the ApPr. This is an important finding that highlights the role of the ApPr as a sensory compartment. In this work we show that when recording from CSFcNs a PKD2L1-mediated current is evoked when uncaging protons on the ApPr, but not when uncaging on the dendrite or the soma. As expected, uncaging a few microns away from the ApPr is inefficient to trigger a response, which confirms the high spatial resolution of the technique^25^. The other experiments presented in **Figure 6** point to the same direction: i. outside-out recordings from the ApPr show PKD2L1-dependent single channel activity; ii. whole-cell recording from isolated ApPrs show single-channel currents that are indistinguishable from those recorded from intact ApPrs; iii. photolysis on isolated ApPr evokes PKD2L1-mediated currents as in intact cells; iv. immunohistochemical experiments show that PKD2L1 expression is higher in the A pPr than in the soma (see also^34^). Although we have not attempted to do outside-out patches from somatic recordings, single channel recordings in this configuration has proved to be difficult by another group^14^. It is interesting to analyse this anatomical segregation of the receptors in the context of previous electron microscopy data, which shows that in the intact spinal cord a tight cytoskeleton ring (mainly adherens and tight junctions) formed by the ependymal cells^35,36^ isolates the ApPr from the rest of the spinal parenchyma. Likewise, we have recently shown that ApPrs have distinct histological features: they are stabilized by a drebrin-rich, specialized actin cytoskeleton and are enriched with non-muscular myosin IIB^37^. Interestingly, it has been shown that the ionic composition of the CSF is different from that in the serum^38^. This implies that the ApPr and the soma are exposed to different extracellular solutions, with a potential impact on the equilibrium potentials of the main ions involved in the CSFcNs electrical response. Although in our slice preparation the interphase separating the CSF from the spinal cord parenchyma is lost, it makes sense that the ApPrs are immersed in the CSF and physically separated, as they should if they are to sense CSF composition.

The physiological features of PKD2L1 channels need also to be taken into account to fully understand the role of CSFcNs as pH sensors: i. PKD2L1 channels are extremely sensitive to proton concentration in the physiological range (**Figure 3**). This has been quantified by González-Perrett et al.^23^, who showed that the equilibrium constant for the related PKD2 channel expressed in lipid bilayers is 6.4 (see also^24^). At this pH value the open probability of the channel is half of its maximum. We performed the same analysis (**Figure 3Ce**) and obtained a value close to 8.0.

Interestingly, the holding current can also be used as a proxy of the channel sensitivity to pH changes (**Figure 3Cf**). The fact that physiological pH indicates the half-maximum of the activation state of the channel is probably the most important biophysical characteristic of PKD2L1 channels in the context of CSFcN physiology; ii. PKD2L1 channels are able to sense deviations of the pH in both directions: an increase in proton concentration (acidification) decreases the channel open probability, while a decrease in proton concentration (alkalinisation) increases the channel open probability. Enhanced excitability of CSFcNs resulting from increased PKD2L1 activity due to alkalinisation has been shown in mice^15^ and lamprey^13^, and has been attributed to augmented phasic activity of the channel. Remarkably, the fact that a decrease in channel activity can also provide meaningful information to the cell by inducing a strong hyperpolarization (**Figure 3B**) has not been described before. This exquisite sensitivity of the cell’s RMP to small changes in the membrane current can be explained by the high input resistance of the ApPr, which we have quantified here by performing direct recordings from that compartment; iii. finally, the physiological effects of PKD2L1 channels are achieved in part through the modulation of a sustained current.

Based on cryo-EM data, Su et al. have attempted to explain the structural mechanisms of the pH sensitivity of the PKD2L1 channel^39^. In their work, the authors compare PKD2L1 channels with another pH-sensitive member of the TRP family, TRPML3, who is also pH sensitive and whose structures at neutral (7.4) and acidic (4.8) pH are known^40^. The luminal loops of PKD2L1 share sequence similarities with those of the TRPML3 channel, and notably the presence of polar residues that may translate the binding of the protons into a conformational change that closes the channel.

### The sustained current

In GABAergic systems, two different types of GABA_A_ receptors mediate phasic and tonic currents: one depends on the activation of low-affinity receptors by synaptically released GABA, while the other depends on the activation of high-affinity GABA_A_ receptors by low concentrations of ambient GABA^41^. In this work we describe phasic, single-channel currents that depend on the spontaneous opening of PKD2L1 channels, and a type of sustained current that was made evident when PKD2L1 channels were blocked with dibucaine. The following experimental evidence indicates that both the phasic, single channel events, and the sustained current depend on the same channel: first, the effect of dibucaine persists when blocking voltage-dependent and ligand-gated channels that may also contribute to the appearance of a sustained current; second, the effect of dibucaine was still present when recording CSFcNs with an IS containing a high concentration of BAPTA; third, the sustained current was modulated by other exogenous (calmidazolium) and endogenous (pH) modulators that are known to affect PKD2L1 channels. These experiments suggest that, apart from the closed and open states of the channel, some other conducting state of the channel may exist as well. Alternatively, very short opening periods or low amplitude of the channels, not seen in the recordings but nevertheless present and blocked by dibucaine, could also explain the sustained current. The assessment between these two putative mechanisms is beyond the scope of this work.

The magnitude of the sustained current compared to the spontaneous, phasic currents, is substantial. The average single channel current measured in this work is ≈ -16 pA and the apparent po1 is ≈ 0.04, which determines a mean phasic current of 16 pA × 0.04 = 0.64 pA. On the other hand, the average sustained current (**Figure 2Ca**) is ≈ 8 pA. Altogether, the percentage of sustained and phasic currents are roughly 90 % and 10 %, respectively, which highlights the importance of the sustained current in the physiology of CSFcNs.

### The photolysis-evoked PKD2L1 recovery current

It has been reported previously that PKD2L1 channels are responsible for a recovery response when challenged by acidic solutions^24,29,30,42^. A first indication of the presence of this recovery response in CSFcNs was provided by Orts-del’Immagine^14^, who reported in some of the neurons recorded in I-clamp an increase in the frequency of depolarising activity after pressure applying a pH 2.8 solution. In this work we report for the first time the presence of this recovery response by using laser photolysis of protons and further show that the current is only evoked when uncaging at the ApPr, indicating that functional PKD2L1 channels are predominantly located there (see above); we call this current the “photolysis-evoked PKD2L1 current”. In the published literature the recovery response, which has been called “off-current”, appears with large-scale acidifications, in the order of several pH units. In this paper we have likewise induced substantial pH changes (to 3 or 3.5; see **Methods**) but have not attempted to quantify the pH dependence of the recovery response. It is nevertheless interesting to highlight the fact that the recovery response does not require long-lasting changes in the extracellular pH to develop. Instead, very brief pH changes are enough, which opens the possibility that the ApPrs may also respond to sudden ACSF acidifications, as may occur during neurosecretion^43^. It would be interesting in the future to explore with photolysis the concentration and time dependence of the recovery current (for example, smaller pH changes but longer pulses) in order to assess its physiological significance. It is also important to take into account that in the zebrafish, PKD2L1 activity is highly dependent of the tension level of the Reissner fiber^44^. In rodents, it remains to be established whether the Reissner fiber plays any role in the activity of CSFcNs (which is not conserved in the acute brain slices used here).

### The activation of PKD2L1 channels in the physiological and pathophysiological context

We have so far discussed how PKD2L1 channels make CSFcNs able to respond to increases and decreases in proton concentration (acidification and alkalinisation, respectively). But in which physiological context is CSFcN activity affected by pH changes? Although CSF pH is tightly regulated, it may change during periods of intense physical activity. This may impact on the activity of CSFcNs that, trough their synaptic connections with neurons of locomotor CPGs in the spinal cord, may adapt motor output in order to help regulate pH. The same type of feedback mechanism to adapt motor output may by implemented in pathophysiological conditions, such as infections, stroke or spinal cord injury, where the CSF pH changes may be more important.

The “recovery” current, on the other hand, is triggered after short and large acidifications (pH ≈ 3.5), as the ones evoked here with laser photolysis. The presence of this current is a fundamental piece of evidence that confirms that functional PKD2L1 channels are segregated to the ApPr, as it does not appear when uncaging in the other neuronal compartments (soma or dendrite). However, it remains to be established whether the recovery current has any physiological meaning. Although very short pH transients are difficult to measure, they do happen in some physiological conditions, for example during transmitter release, when the very acidic synaptic vesicles fuse with the plasma membrane^45^ in a very restricted volume (the synaptic cleft). In these conditions, where the pH changes phasically within the millisecond time-range in a very tiny volume, the appearance of the above-mentioned “recovery current” may happen. It was recently shown by Prendergast and collaborators^43^ that in some situations of bacterial infections, CSFcNs may release different molecules to the central canal. This creates the perfect environment for the appearance of the PKD2L1-dependent recovery current.

### The electrotonic coupling between the ApPrs and the soma

It was shown previously that the activation of a single PKD2L1 channel can induce spiking ^15^. As PKD2L1 channels are enriched in the ApPr, where we found no evidence for sodium channels, the voltage changes produced by their activation need to propagate very efficiently to the action potential triggering zone in order to initiate spiking. Here, we performed for the first time paired ApPr and somatic recordings in order to measure directly the amount of coupling between these 2 compartments. We show that the coupling for spontaneous events is around 0.88 and even higher for square current pulses (the DC CC), in agreement with what has been previously shown with modeling by Orts-Del’immagine and collaborators, who predicted a 7% attenuation for somatically recorded voltage changes originating in the ApPr. The average conductance between the 2 compartments, on the other hand, is around 13 nS. The high single channel conductance of PKD2L1 channels, together with the high input resistance of the ApPr and its strong electrotonic coupling with the soma, all contribute to the efficient passive propagation of signals originating in the ApPr.

### PKD2L1 downstream signaling mechanisms

The anatomical segregation of the ApPr within the cc and the biophysical properties of PKD2L1 channels suggest that the ApPr compartment may have more complex functions rather than simply affecting CSFcN firing. Indeed, it seems that calcium flowing into the cell through PKD2L1 channels is a key regulator of the ApPr physiology: on the one hand, PKD2L1 channels are highly calcium permeable and at the same time, they are inhibited by intracellular calcium^19^. Also, ultrastructural data indicate that the ApPr is rich in mithocondria and tubo-vesicular structures resembling the Golgi apparatus^35,36^, which are intracellular organelles that contribute to calcium homeostasis. Altogether, this evidence suggests that intraApPr calcium concentration needs to be finely regulated, both in space and in time, in order for the ApPr to fulfill its physiological roles. Based on what has been published in the literature, we can speculate that these calcium signals can be decoded by several systems: i) calcium can act as the second messenger linking the activation of the multimodal PKD2L1 channels to changes in CSFcNs excitability, which in turn regulate spinal neuronal networks controlling locomotor activity; ii) calcium could initiate neurosecretion of different molecules from the ApPr to the central canal (as has been proposed by the Wyart group in the zebrafish in the context of bacterial infections^43^); iii) calcium could activate the Hedgehog signaling pathways^46^; iv) calcium could modulate CSF flow directly (by modulating filopodial activity^37^) or indirectly (by modulating ependymal cells ciliary activity through paracrine interactions). Resolving these downstream pathways is essential to fully understand the role of CSFcNs as integrators of CSF homeostasis.

### The involvement of ASIC

ASICs, also known as proton-gated channels, play important physiological and pathological roles in the nervous system^45,47^. In CSFcNs, ASICs have been reported as mediating part of the response to acidification^13–15,31^. In this study, we observed in a few cells a rapidly desensitizing inward current when pressure applying a pH 6.5 solution, and occasionally an inward current with similar characteristics when uncaging either MNI-Glu or MNI-γLGG at the soma (**Supplementary Figure 2B**). However, we never saw such an inward current when photolysing at the ApPr, and the photolysis-evoked PKD2L1 current evoked by proton release on the ApPr was not affected in the presence of ASIC channel blockers (**Figure 5E**). As ASICs are widely expressed in many brain areas and neuronal compartments^33^, the most parsimonious explanation for these experimental results is that the ASIC-dependent current recorded in CSFcNs results from the activation of somatic ASICs, but that those channels are not present in the ApPr. The picture that emerges from our data, together with results published previously by other groups, suggests that the pH sensitivity of CSFcNs is thus a complex response that depends on different channels localized in different neuronal compartments. ASICs may be suited to sense pH changes in the spinal cord parenchyma, next to CSFcN soma, while PKD2L1 channels may have adapted to sense pH changes in the CSF. In other words, CSFcNs pH sensitivity is also a compartmentalized function. It remains to be established which are the exact conditions (in terms of absolute pH changes and kinetics) that activate each of the channels and how the cell integrates these signals in order to produce a meaningful response.

## Materials and methods

### Transgenic mice

Gata3^eGFP^ mice were generated using bacterial artificial chromosome (BAC) recombination as described^48^. The eGFP gene was inserted by deleting 286 bp of coding sequences from the first exon. The sequences flanking the insertions were: 5’agccgaggac-eGFP–cgtggaccca3’. Four Gata3-eGFP founder lines were generated that expressed eGFP in an identical manner to GATA3 in the spinal cord. One of these lines was selected for this study. Animals of either sex were maintained in a 12 hs. light/dark cycle with food and water *ad libitum*, humidity between 50 and 60%.

### Slice preparation for electrophysiological experiments

Spinal cord slices were obtained from Gata3^eGFP^ mice aged 30 to 45 days old. Slices were prepared as previously described^3^. Briefly, mice were decapitated under isoflurane anesthesia following approved ethical procedures (CEUA 001/01/2022a) and the spinal cord dissected in an ice-cold artificial ACSF of the following composition (in mM): 101 NaCl, 3.8 KCl, 1.3 MgSO_4_.7H_2_O, 1.2 KH_2_PO_4_, 10 HEPES, 25 Glucose, 1 CaCl_2_, 18.7 MgCl_2_, osmolarity 300 mOsm/kg H_2_O and pH adjusted to 7.4. The spinal cord was included in a 4% low melting point agar in order to glue it in the desired orientation. 300 µm thick lumbar spinal cord slices were obtained following 2 dissecting planes, either transverse or with a 45 degrees angle. The latter orientation was used because we found that the probability of obtaining superficial, and hence easier to patch, ApPrs, was higher. Also, we found that the chances of obtaining “isolated” ApPrs (see **Figure 6B**) was also higher in the oblique preparation.

### Electrophysiological recordings and epifluorescence

CSFcNs were recorded with the patch-clamp technique^49^ at near physiological temperature (34 ± 1 °C, Peltier system; Luigs & Neurmann) on an upright Olympus microscope (BX51WI) equipped with a 60×, 1.0 numerical aperture objective. Once in the recording chamber, slices were continuously perfused with the following extracellular solution (in mM): 115 NaCl, 2.5 KCl, 1.3 NaH_2_PO_4_, 26 NaHCO_3_, 25 glucose, 5 NaPyruvate, 2 CaCl_2_.2H_2_O and 1 MgCl_2_.6H_2_O, osmolarity 300 mOsm/kg H_2_O and pH 7.4 with the continuous bubbling of a mixture of 95% O_2_ and 5% CO_2_. Recordings were performed with a KGluconate-based intracellular solution of the following composition (in mM): 165 KGluconate, 10 HEPES, 1 EGTA, 0.1 CaCl_2_, 4.6 MgCl_2_, 4 Na_2_ATP, 0.4 NaGTP and 0.04 Alexa 594, pH 7.3, and osmolarity 300 mOsm/kg H_2_O. With this solution, pipette resistance was 6 to 7 MΩ for somatic and 10 to 11 MΩ for ApPr recordings; series resistance, on the other hand, was monitored during the whole experiment, but not compensated for. Recordings were performed with a HEKA amplifier (EPC10 USB) and the software Patchmaster either in voltage-clamp (-60 mV holding potential) or I-clamp. Recordings were acquired at a sampling rate of 20 kHz and low-pass filtered at 2.9 kHz. Recording and puffing pipettes were positioned in the slice with Luigs & Neumann micromanipulators. Epifluorescence excitation was by light-emitting diodes in a dual lamp-house (Optoled; Cairn Research). EGFP was excited at 470/40 nm and Alexa 594 at 572/35 nm. Fluorescence emission at 520/40 nm and 630/60 nm was detected with an EM CCD camera (Andor Ixon) and filters from Chroma Corporation (Vermont, USA).

### Paired recordings apical process – soma

Both the ApPr and the soma of a single neuron were simultaneously patched following usual procedures. No Alexa 594 was included in the IS. In order to maximize the chances of getting a paired recording, the patching sequence was as follows: somatic cell-attached, ApPr cell-attached, somatic break-in and ApPr break-in. The steady-state coupling coefficient (DC CC) was measured in I-clamp by injecting rectangular current pulses either in the ApPr or in the soma. Plots of the voltage changes in the non-pulsed vs the pulsed compartments were constructed, and the DC CC was the slope of the linear regression (**Figure 7A**). The coupling coefficient for spontaneous synaptic potentials (ssp CC) was measured by calculating the ratio between the spontaneous potentials recorded in the soma and those recorded in the ApPr (average somatic ssp / average ApPr ssp calculated from at least 10 individual ssps). The conductance value between the 2 compartments, that we call here the coupling conductance, was measured in voltage-clamp by pulsing either of the 2 compartments (from -110 to -20 mV in 10 mV steps, from a holding potential of -60 mV) and constructing an I/V curve from the current recovered in the non-pulsed compartment. In these conditions, the coupling conductance was the slope of the linear fit of this I/V curve.

### Photolysis

Protons were photo-released either from MNI-Glutmate (4-Methoxy-7-nitroindoli-inyl-caged-L-glutamate) or MNI-γLGG (4-Methoxy-7-nitroindoli-inyl-caged-γ-L-glutamyl-glycine) with a 405-nm diode laser (Obis, Coherent, USA) that was focused through the ×60/NA1.0 water immersion objective. The cages were diluted in the recording ACSF to a final concentration of 0.5 or 1 mM and either pressure-applied with a Picospritzer III (Parker) for at least 10 seconds before applying the light pulses, or bath applied. Photolysis light pulses had durations of 0.5 to 1 ms and 1 to 5 mW laser power, and were repeated every 5 seconds. The power of the laser-pulse was monitored on a calibrated photodiode. The laser spot was fixed, and the photolysis location chosen by positioning the slice on the region of interest. Before each experiment, the exact location of the laser spot was verified by monitoring it with a fluorescent solution.

In order to measure the distribution of the laser light intensity, the focused 405 nm laser light was reflected back to the camera by a mirror that was placed at the location of the specimen ^25,50,51^. In these conditions, the lateral, x-y, dimensions of the focused beam were close to 1 µm (1/e^2^ diameter 1.1 µm) and the axial, x-z, dimensions close to 5 µm (1/e^2^ diameter 4.8 µm).

Under the conditions mentioned in the preceding paragraph, photolysis could be repeated multiple times (at least 10) in the same spot without any indication of run-down, adaptation or desensitization of the receptors.

### Chemicals

Salts were either from Sigma-Aldrich or Carlo Erba. TTX (tetrodotoxin; 0.4 µM), TEA (tetraethylammonium; 1.5 mM), 4AP (4 aminopyridine; 1 mM), NBQX (2,3-Dihydroxy-6-nitro-7-sulfamoyl-benzo[f]quinoxaline; 10 µM), D-APV (2-Amino-5-phosphonovaleric acid; 50 µM) and gabazine (10 µM) were from Tocris. MNI-glutamate (1 mM) was from HelloBio and MNI-γLGG (1 mM) was a generous gift from Dr. Céline Auger (Sppin, CNRS UMR8003). Dibucaine hydrochloride (100 to 200 µM) and calmidazolium chloride (20 µM) were from Sigma-Aldrich. Psalmotoxin 1 (100 nM), APETx2 (100 nM) and TTA-P2 (20 µM) were from Alomone Lab.

### Immunohistochemistry

Animals were anesthetized with ketamine (100 mg/kg, i.p.), xylacine (10 mg/kg, i.p.), and diazepam (5 mg/kg, i.p.) and fixed by intracardiac perfusion with 4% PFA in 0.1 M PB. The spinal cord was sectioned with a vibrating microtome (50 - 70 µm thick), placed in PBS with 0.5% BSA for 30 min, and then incubated with the primary mouse anti-PKD2L1 antibody (Millipore-Sigma AB9084, 1:500) in PBS with 0.3% Triton X-100 (Sigma Millipore). Sections were then incubated in the secondary donkey anti-mouse Alexa-647 antibody (Thermo-Fisher A-21235, 1:1000) and mounted in 70% (v/v) glycerol pH 8.8. Nuclei were stained with 1µg/ml Hoechst 33342 (Thermo-Fisher).

### Confocal microscopy and image processing

Line selections were manually drawn through either the soma or ApPr of GFP+ cells on single deconvolved confocal planes and fluorescence intensity values per pixel along the selection were obtained using FIJI/ImageJ^52^. Descriptive statistical values were used to build the average and SD plots against position along the selection (x).

Images were acquired on a Zeiss LSM800 confocal microscope with a 63× oil immersion lens (NA

= 1.4), pinhole set at 0.8 Airy units. Sampling interval was set to an oversample density according to the Nyquist criterion. Z-stacks were deconvolved with Huygens Essential 4.5 (Scientific Volume Imaging B.V., Hilversum, Netherlands) using an experimental PSF, SNR = 20, using an iterative Classic Maximum Likelihood Estimation (CMLE) algorithm set to a maximum of 40 iterations, with a quality threshold of 0.05. Composite image assembly and further processing (brightness/contrast adjustment), when needed, was performed using FIJI/ImageJ.

### Analysis

Electrophysiological analysis was performed with Igor Pro (Wavemetrics) using Neuromatic^53^, TaroTools (https://sites.google.com/site/tarotoolsmember/?authuser=1) and custom routines. Single-channel currents were analyzed with the software Nest-O-Patch (NOP; https://sourceforge.net/ projects/nestopatch/). The currents were filtered with the built-in filter of the software (cutt-off filter of 1.4 KHz) and the single-channel current amplitude calculated from a multigaussian fit adjusted to the whole-cell current histogram. Apparent closed and open probabilities were calculated from the idealized current traces. The obtained values were then imported to IgorPro (8.0 or 9.0, Wavemetrics, Lake Oswego) for further analysis.

As the majority of the recordings presented in the manuscript are not long enough to calculate a meaningful po, we have calculated 2 other parameters from our recordings: n_max_, which corresponds to the maximum number of channels that open simultaneously during a 500 ms time window, and t_open_, which corresponds to the total open time of a single channel during the same 500 ms time window. It is reasonable to think that the total amount of channels does not change during the electrophysiological experiments, so n_max_ and t_open_ reflect an indirect measure of po.

In **Figure 3Ce**, where the activity of the channel was measured as a function of different pHs, po1 was calculated during time periods of 2 seconds. Then, the values obtained were normalized to the control value (pH 7.4). We call this the normalized apparent po1.

Current charge was calculated as the integral of the whole-cell recordings after baseline subtraction. The number of recordings averaged in each condition is indicated in the corresponding figure legend.

Holding current in different conditions was calculated as follows: histograms were constructed from current recordings during 2 to 3 seconds time epochs. Then, the first peak of the resulting histogram, which corresponds to the baseline current, was fitted with a Gaussian function (IgorPro built-in function); the mean value of the fitting corresponds to the sustained current (see **Figure 2Bb** for an example). In order to show visually the sustained current change in different conditions, current traces were averaged over 100 ms epochs, like what has been done for the analysis of GABA_A_-mediated tonic currents^54^. For the analysis of resting membrane potential in different conditions, average Vm values were calculated from 1 second long time periods.

Input resistance was measured in V-clamp by applying voltage steps from -110 or -100 to -60 mV in 10 mV increments. From these pulses the steady-state current was measured and the input resistance calculated from the slope of the I/V curve. At these potentials no voltage-dependent conductances were activated.

### Estimation of the pH drop induced by photolysis

The photolysis of MNI-based compounds leads to proton release. The pH drop induced by the photolysis reaction depends on how many protons are released, on the speed of the buffering system and on the diffusion of molecules in and out of the illuminated volume. In our experiments we used caged-compounds concentrations between 0.5 and 1 mM. Based on our previous callibrations^25^ that indicate that the photolysis is complete with the energy used here (≈ 2 mW for 1 ms), we expect an added charge of 0.5 to 1 mM protons in the illuminated volume, corresponding to a drop of pH from 7.4 to ≈ 3-3.5 pH units. This acid pH is quickly buffered by bicarbonate. Given that the speed of protonation of bicarbonate is extremely fast^55–57^, the pH returns to 7.4 in less than 1 ms. This indicates that the photolysis-evoked current (**Figures 4**, **5** and **6**) has very similar characteristics to the PKD2L1-mediated current that has been described in expression systems: it is a current produced by the opening of the channel when the acidic stimulus is removed. The fact that this recovery current is so clearly seen when performing photolysis experiments and not with other methods, like pressure-application, likely reflects the superior temporal resolution of photouncaging. This approach enables exceptionally rapid pH jumps. Similarly, it has been shown by Hu et al^58^ that using fast application methods is a necessary condition to induce a type of response, that the authors called Ca^++^ spike, when applying extracellular Ca^++^ stimuli.

### Statistics

Data are presented as mean ± SD. Statistical significance was assessed with the Wilcoxon signed-rank test for paired and the Wilcoxon–Mann–Whitney test for non-paired data. The difference between groups was considered significant when p < 0.05; when significant, the exact p value is indicated in each figure and figure legends.

## Acknowledgments.

This work was supported by a Agencia Nacional de Investigación e Innovación grant (number FCE_1_2021_1_166464) to Federico F. Trigo and a grant from the Wings For Life Spinal Cord Research Foundation (number WFL-UY-13/23 #290) to Raúl R. Russo. Magdalena Vitar was supported by ANII through a Master Student fellowship. We thank Victoria Falco Pastorino and María Gabriela Fabbiani for their help with the maintenance of the Gata3^eGFP^ colony, Céline Auger for the generous gift of MNI-γLGG, John ET Corrie for initial discussions on the proton photolysis and Alain Marty for the critical reading of the manuscript. We would also like to thank Gonzalo Budelli and Gonzalo Pizarro for helpful discussions.

---

**Supplementary Fig. 1.**
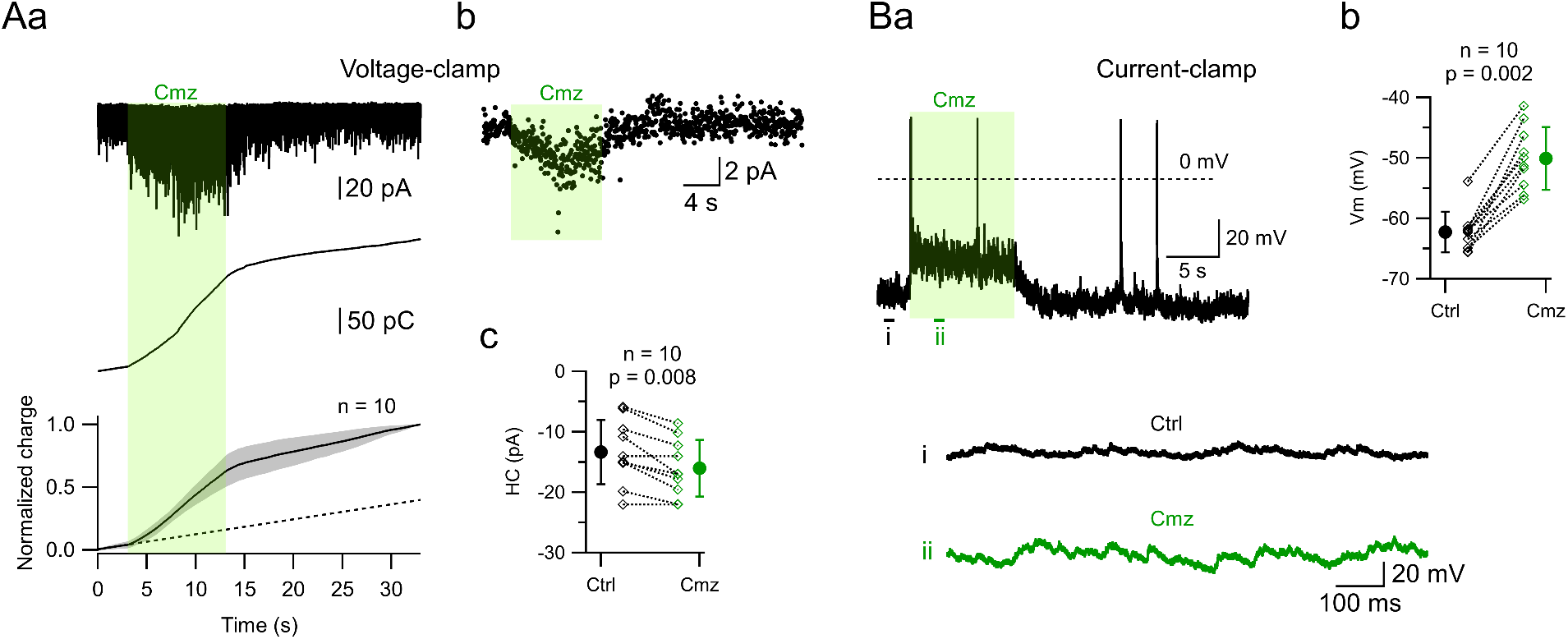
Calmidazolium effect on PKD2L1-mediated tonic and sustained currents. **Aa. Top**. Spontaneous PKD2L1 channel activity recorded in a CSFcN at a -60 mV holding potential. The recording lasted 35 s, and calmidazolium was pressure-applied at a concentration of 10 µM during 10 s (starting at 3 s, green area). **Middle.** Membrane charge calculated from the recording on the top. **Bottom.** Mean membrane charge ± SD calculated from 10 neurons tested in the same conditions as the cell shown in the top panel. The dotted line corresponds to a linear fit to the control period (first 3 seconds of the recording). The application of calmidazolium produces an increase in the slope of the membrane charge. **Ab.** Holding current from the neuron shown in “**a**”. **Ac.** Effect of calmidazolium application on the holding current: -13.3 ± 5.3 pA in control conditions vs -16.0 ± 4.7 pA in calmidazolium (n = 10, p = 0.008). **Ba.** Spontaneous activity of the CSFcN shown in **A**, recorded in I-clamp. Calmidazolium application is indicated with the green area. The bottom traces show segments **i** and **ii** with different time and amplitude scales in order to see the increase in spontaneous activity during calmidazolium. **Bb.** Effect of calmidazolium application on the resting membrane potential. Notice the shift from -62.3 ± 3.3 in control conditions vs -50.0 ± 5.2 mV in the presence of calmidazolium (n = 10, p = 0.002). In **Ac** and **Bb**, diamonds correspond to individual neurons and circles to the AV ± SD. Cmz: calmidazolium. In **Ac** and **Bb**, statistical comparison between groups was performed with a Wilcoxon signed-rank test.

**Supplementary Fig. 2.**
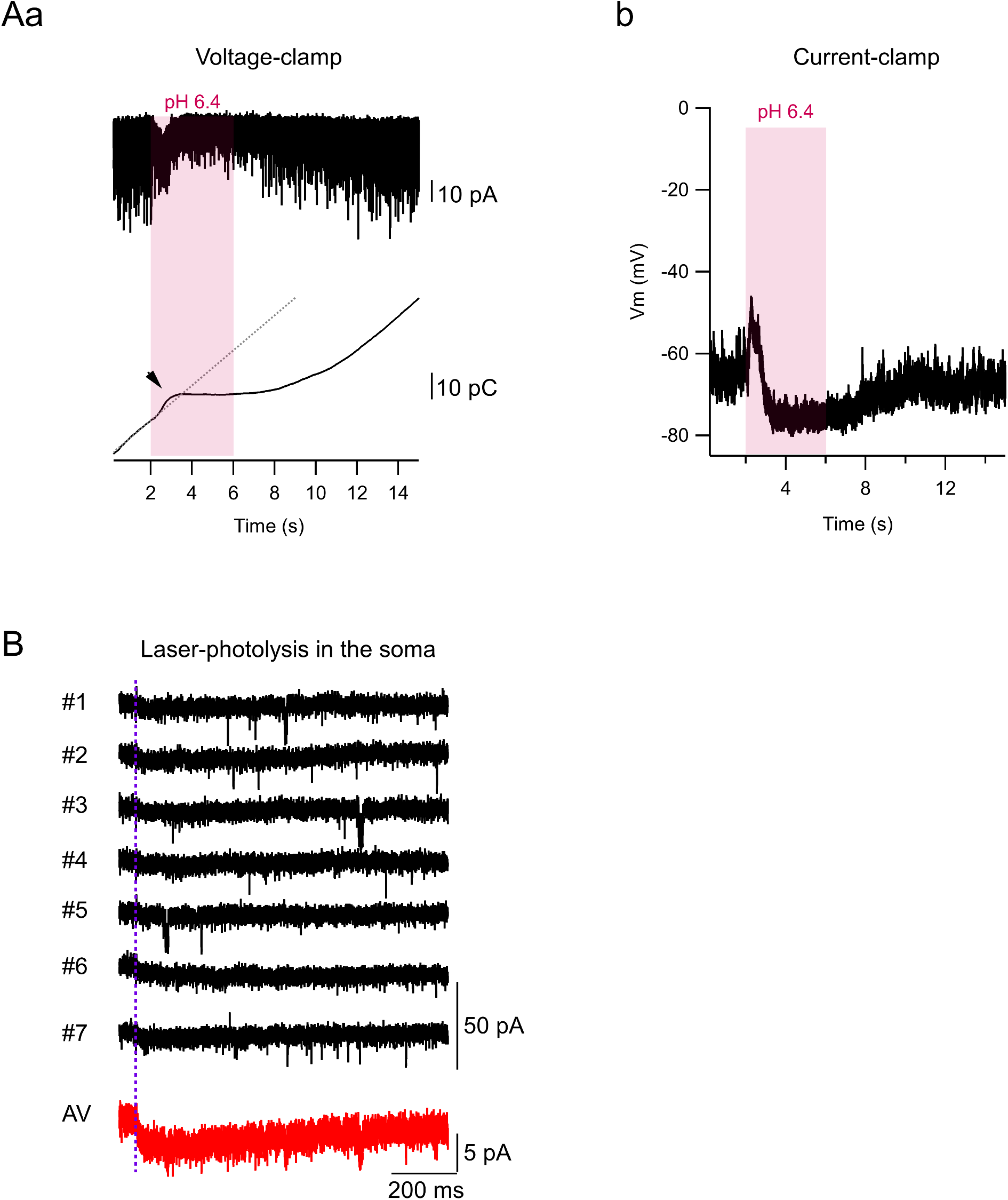
Puffing acidic solutions and proton photolysis in the soma can induce ASIC-mediated currents. **Aa. Top.** Spontaneous PKD2L1 channel activity recorded in a CSFcN at a -60 mV holding potential. A pH 6.5 solution was pressure-applied during 4 s (starting at 2 s, magenta area). **Bottom.** Membrane charge calculated from the recording on the top. The dotted line corresponds to a linear fit during the control period (first 2 seconds of the recording). The application of the acidic solution produces a short-lasting inward current that is probably due to the activation of somatic ASIC channels. This is manifested as a sudden increase in the membrane charge (black arrowhead) that is followed by a subsequent decrease. **Ab.** Same experiment as in **a**, under I-clamp. The application of the acidic solution produces a short-lasting depolarization that is probably due to the activation of somatic ASIC channels, followed by hyperpolarization. Vm: membrane potential. **B.** Spontaneous activity recorded in a CSFcN upon the somatic uncaging of MNI-Glutamate in the continuous presence of NBQX. The black traces show individual repetitions (7 uncagings at 10 sec intervals) and the red trace the corresponding average (AV). Vertical magenta arrowhead and line show the timing of the laser pulse (1 ms, 3 mW). Somatic photolysis does not evoke any PKD2L1-dependent activity, but does produce a small inward current that is probably mediated by ASIC channels.

**Supplementary Fig. 3.**
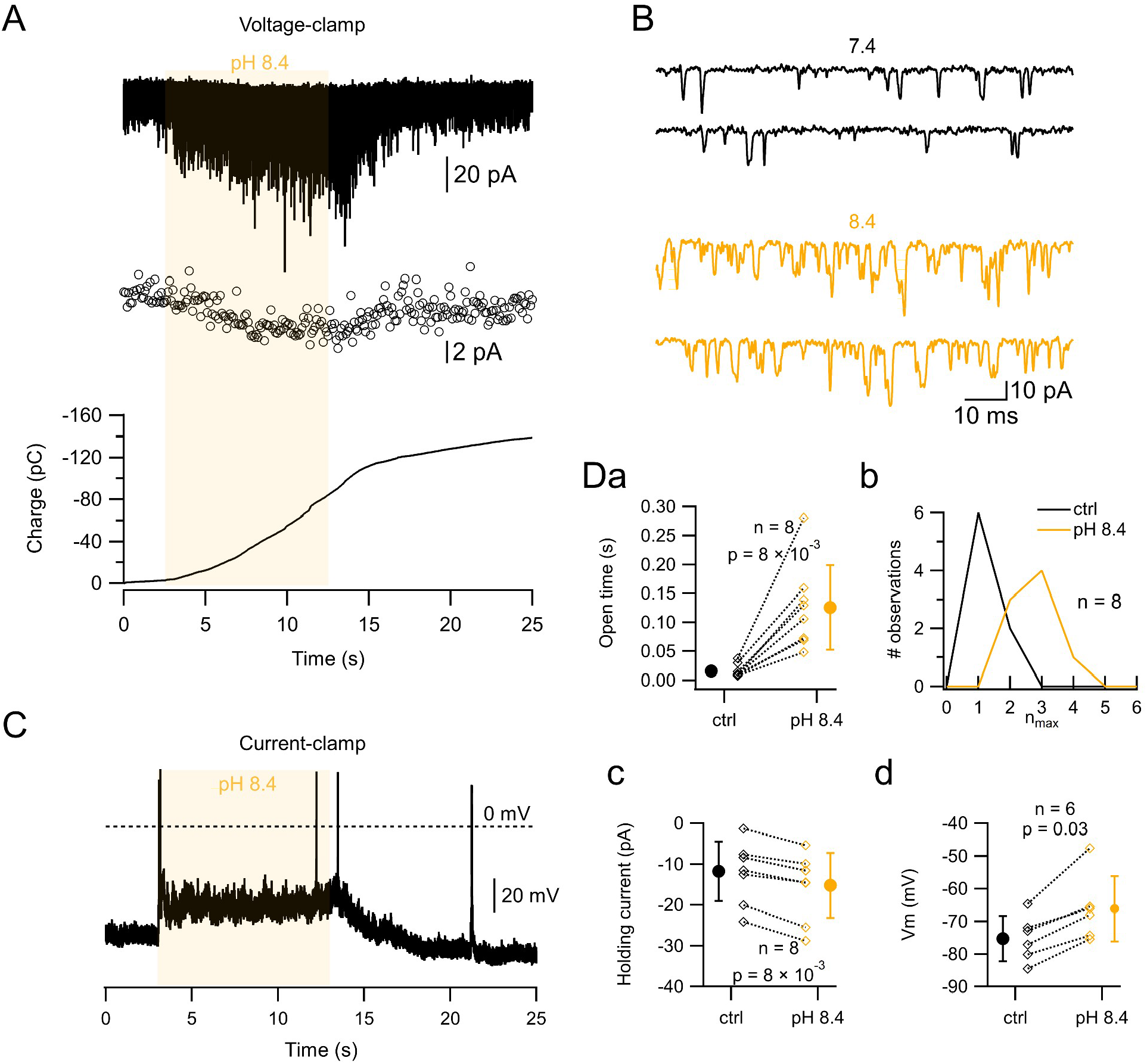
Effect of alkaline ACSF on PKD2L1-mediated currents. **Aa. Top**. Spontaneous PKD2L1 channel activity recorded in a CSFcN at a -60 mV holding potential. A pH 8.4 solution was pressure-applied during 10 s (starting at 3 s, yellow area). **Middle.** Holding current calculated from the recording shown on the top. **Bottom.** Membrane charge calculated from the recording shown on the top. **B.** Chosen segments of the recording shown in **A** with different scales. **C.** Same experiment as in **A**, but the spontaneous activity was recorded in I-clamp. The application of the basic solution produces a depolarization of the RMP (from -64.5 to -47.6 mV in this example). **Da.** Effect of the pH 8.4 solution on the total open time of a single channel during a 500 ms time window. The mean single channel open time increased from 16 ± 11 (ctrl) to 126 ± 73 ms (pH 8.4; n = 8, p = 0.008). **Db.** Histograms of the n_max_ values calculated during 500 ms time windows for spontaneous recordings (black curve) and during the application of a pH 8.4 solution (magenta curve). The mean n_max_ increased 1.25 ± 0.46 (ctrl) to 2.75 ± 0.70 (pH 8.4; n = 8, p = 0.016). A two sample Kolmogorov-Smirnov test also indicated a significant difference between the 2 distributions (p = 0.011). **Dc.** The application of a pH 8.4 solution produced a shift of the holding current from -11.7 ± 7.3 to -15.2 ± 7.9 pA (n = 8, p = 0.0078). **Dd.** A pH 8.4 solution induced a shift of the resting membrane potential from -75.2 ± 7.0 vs -66.2 ± 10 mV (n = 6, p = 0.03). In **D**, statistical comparison between groups was performed with a Wilco

## Notes

### Competing Interest Statement

The authors have declared no competing interest.

### Summary of Updates

Minor change in the Title (the word exclusive was deleted). Minor changes in the article ("tonic" has been replaced by "sustained" and "off-current" has been replaced by "recovery current". Supplementary Figure 4 has been moved as a main Figure (Figure 5) and Figures 5 and 6 renamed 6 and 7, respectively.

## References

(1) Wyart, C.; Carbo-Tano, M.; Cantaut-Belarif, Y.; Orts-Del’Immagine, A.; Böhm, U. L. Cerebrospinal Fluid-Contacting Neurons: Multimodal Cells with Diverse Roles in the CNS. Nat Rev Neurosci 2023, 24 (9), 540–556. 10.1038/s41583-023-00723-8.

(2) Wang, R. L.; Chang, R. B. The Coding Logic of Interoception. Annual Review of Physiology 2024, 86 (Volume 86, 2024), 301–327. 10.1146/annurev-physiol-042222-023455.

(3) Marichal, N.; García, G.; Radmilovich, M.; Trujillo-Cenóz, O.; Russo, R. E. Enigmatic Central Canal Contacting Cells: Immature Neurons in “Standby Mode”? J. Neurosci. 2009, 29 (32), 10010–10024. 10.1523/JNEUROSCI.6183-08.2009.

(4) Orts-Del’Immagine, A.; Kastner, A.; Tillement, V.; Tardivel, C.; Trouslard, J.; Wanaverbecq, N. Morphology, Distribution and Phenotype of Polycystin Kidney Disease 2-like 1-Positive Cerebrospinal Fluid Contacting Neurons in the Brainstem of Adult Mice. PLoS One 2014, 9 (2), e87748. 10.1371/journal.pone.0087748.

(5) Hubbard, J. M.; Böhm, U. L.; Prendergast, A.; Tseng, P.-E. B.; Newman, M.; Stokes, C.; Wyart, C. Intraspinal Sensory Neurons Provide Powerful Inhibition to Motor Circuits Ensuring Postural Control during Locomotion. Curr Biol 2016, 26 (21), 2841–2853. 10.1016/j.cub.2016.08.026.

(6) Gerstmann, K.; Jurčić, N.; Blasco, E.; Kunz, S.; de Almeida Sassi, F.; Wanaverbecq, N.; Zampieri, N. The Role of Intraspinal Sensory Neurons in the Control of Quadrupedal Locomotion. Curr Biol 2022, 32 (11), 2442–2453.e4. 10.1016/j.cub.2022.04.019.

(7) Nakamura, Y.; Kurabe, M.; Matsumoto, M.; Sato, T.; Miyashita, S.; Hoshina, K.; Kamiya, Y.; Tainaka, K.; Matsuzawa, H.; Ohno, N.; Ueno, M. Cerebrospinal Fluid-Contacting Neuron Tracing Reveals Structural and Functional Connectivity for Locomotion in the Mouse Spinal Cord. Elife 2023, 12, e83108. 10.7554/eLife.83108.

(8) Fidelin, K.; Djenoune, L.; Stokes, C.; Prendergast, A.; Gomez, J.; Baradel, A.; Del Bene, F.; Wyart, C. State-Dependent Modulation of Locomotion by GABAergic Spinal Sensory Neurons. Curr Biol 2015, 25 (23), 3035–3047. 10.1016/j.cub.2015.09.070.

(9) Wu, M.-Y.; Carbó-Tano, M.; Mirat, O.; Lejeune, F.-X.; Roussel, J.; Quan, F.; Fidelin, K.; Wyart, C. Spinal Sensory Neurons Project onto Hindbrain to Stabilize Posture and Enhance Locomotor Speed. bioRxiv 2021, 2021.03.16.435696. 10.1101/2021.03.16.435696.

(10) Jalalvand, E.; Robertson, B.; Wallén, P.; Grillner, S. Ciliated Neurons Lining the Central Canal Sense Both Fluid Movement and pH through ASIC3. Nat Commun 2016, 7, 10002. 10.1038/ncomms10002.

(11) DeCaen, P. G.; Delling, M.; Vien, T. N.; Clapham, D. E. Direct Recording and Molecular Identification of the Calcium Channel of Primary Cilia. Nature 2013, 504 (7479), 315–318. 10.1038/nature12832.

(12) Huang, A. L.; Chen, X.; Hoon, M. A.; Chandrashekar, J.; Guo, W.; Tränkner, D.; Ryba, N. J. P.; Zuker, C. S. The Cells and Logic for Mammalian Sour Taste Detection. Nature 2006, 442 (7105), 934–938. 10.1038/nature05084.

(13) Jalalvand, E.; Robertson, B.; Tostivint, H.; Wallén, P.; Grillner, S. The Spinal Cord Has an Intrinsic System for the Control of pH. Curr. Biol. 2016, 26 (10), 1346–1351. 10.1016/j.cub.2016.03.048.

(14) Orts-Del’immagine, A.; Wanaverbecq, N.; Tardivel, C.; Tillement, V.; Dallaporta, M.; Trouslard, J. Properties of Subependymal Cerebrospinal Fluid Contacting Neurones in the Dorsal Vagal Complex of the Mouse Brainstem. J. Physiol. (Lond.) 2012, 590 (16), 3719– 3741. 10.1113/jphysiol.2012.227959.

(15) Orts-Del’Immagine, A.; Seddik, R.; Tell, F.; Airault, C.; Er-Raoui, G.; Najimi, M.; Trouslard, J.; Wanaverbecq, N. A Single Polycystic Kidney Disease 2-like 1 Channel Opening Acts as a Spike Generator in Cerebrospinal Fluid-Contacting Neurons of Adult Mouse Brainstem. Neuropharmacology 2016, 101, 549–565. 10.1016/j.neuropharm.2015.07.030.

(16) Panayi, H.; Panayiotou, E.; Orford, M.; Genethliou, N.; Mean, R.; Lapathitis, G.; Li, S.; Xiang, M.; Kessaris, N.; Richardson, W. D.; Malas, S. Sox1 Is Required for the Specification of a Novel P2-Derived Interneuron Subtype in the Mouse Ventral Spinal Cord. J. Neurosci. 2010, 30 (37), 12274–12280. 10.1523/JNEUROSCI.2402-10.2010.

(17) Petracca, Y. L.; Sartoretti, M. M.; Di Bella, D. J.; Marin-Burgin, A.; Carcagno, A. L.; Schinder, A. F.; Lanuza, G. M. The Late and Dual Origin of Cerebrospinal Fluid-Contacting Neurons in the Mouse Spinal Cord. Development 2016, 143 (5), 880–891. 10.1242/dev.129254.

(18) Sternberg, J. R.; Prendergast, A. E.; Brosse, L.; Cantaut-Belarif, Y.; Thouvenin, O.; Orts-Del’Immagine, A.; Castillo, L.; Djenoune, L.; Kurisu, S.; McDearmid, J. R.; Bardet, P.-L.; Boccara, C.; Okamoto, H.; Delmas, P.; Wyart, C. Pkd2l1 Is Required for Mechanoception in Cerebrospinal Fluid-Contacting Neurons and Maintenance of Spine Curvature. Nat Commun 2018, 9 (1), 3804. 10.1038/s41467-018-06225-x.

(19) DeCaen, P. G.; Liu, X.; Abiria, S.; Clapham, D. E. Atypical Calcium Regulation of the PKD2-L1 Polycystin Ion Channel. Elife 2016, 5, e13413. 10.7554/eLife.13413.

(20) Guadarrama, E.; Vanoye, C. G.; DeCaen, P. G. Defining the Polycystin Pharmacophore Through HTS & Computational Biophysics. bioRxiv 2025, 2025.01.13.632808. 10.1101/2025.01.13.632808.

(21) Russo, R. E.; Fernández, A.; Reali, C.; Radmilovich, M.; Trujillo-Cenóz, O. Functional and Molecular Clues Reveal Precursor-like Cells and Immature Neurones in the Turtle Spinal Cord. J Physiol 2004, 560 (Pt 3), 831–838. 10.1113/jphysiol.2004.072405.

(22) Park, E. Y. J.; Baik, J. Y.; Kwak, M.; So, I. The Role of Calmodulin in Regulating Calcium-Permeable PKD2L1 Channel Activity. Korean J Physiol Pharmacol 2019, 23 (3), 219–227. 10.4196/kjpp.2019.23.3.219.

(23) González-Perrett, S.; Batelli, M.; Kim, K.; Essafi, M.; Timpanaro, G.; Moltabetti, N.; Reisin, I. L.; Arnaout, M. A.; Cantiello, H. F. Voltage Dependence and pH Regulation of Human Polycystin-2-Mediated Cation Channel Activity *. Journal of Biological Chemistry 2002, 277 (28), 24959–24966. 10.1074/jbc.M105084200.

(24) Chen, P.; Wu, J.; Zhao, J.; Wang, P.; Luo, J.; Yang, W.; Liu, X. PKD2L1/PKD1L3 Channel Complex with an Alkali-Activated Mechanism and Calcium-Dependent Inactivation. Eur Biophys J 2015, 44 (6), 483–492. 10.1007/s00249-015-1040-y.

(25) Trigo, F. F.; Corrie, J. E. T.; Ogden, D. Laser Photolysis of Caged Compounds at 405 Nm: Photochemical Advantages, Localisation, Phototoxicity and Methods for Calibration. Journal of Neuroscience Methods 2009, 180 (1), 9–21. 10.1016/j.jneumeth.2009.01.032.

(26) Blanchard, K.; Zorrilla de San Martín, J.; Marty, A.; Llano, I.; Trigo, F. F. Differentially Poised Vesicles Underlie Fast and Slow Components of Release at Single Synapses. J. Gen. Physiol. 2020, 152 (5). 10.1085/jgp.201912523.

(27) Canepari, M.; Nelson, L.; Papageorgiou, G.; Corrie, J. E. T.; Ogden, D. Photochemical and Pharmacological Evaluation of 7-Nitroindolinyl-and 4-Methoxy-7-Nitroindolinyl-Amino Acids as Novel, Fast Caged Neurotransmitters. Journal of Neuroscience Methods 2001, 112 (1), 29–42. 10.1016/S0165-0270(01)00451-4.

(28) Palma-Cerda, F.; Papageorgiou, G.; Barbour, B.; Auger, C.; Ogden, D. Photolysis of a Caged, Fast-Equilibrating Glutamate Receptor Antagonist, MNI-Caged γ-D-Glutamyl-Glycine, to Investigate Transmitter Dynamics and Receptor Properties at Glutamatergic Synapses. Front Cell Neurosci 2018, 12, 465. 10.3389/fncel.2018.00465.

(29) Hussein, S.; Zheng, W.; Dyte, C.; Wang, Q.; Yang, J.; Zhang, F.; Tang, J.; Cao, Y.; Chen, X.-Z. Acid-Induced off-Response of PKD2L1 Channel in Xenopus Oocytes and Its Regulation by Ca2+. Sci Rep 2015, 5, 15752. 10.1038/srep15752.

(30) Inada, H.; Kawabata, F.; Ishimaru, Y.; Fushiki, T.; Matsunami, H.; Tominaga, M. Off-response Property of an Acid-activated Cation Channel Complex PKD1L3–PKD2L1. EMBO reports 2008, 9 (7), 690–697. 10.1038/embor.2008.89.

(31) Jalalvand, E.; Robertson, B.; Tostivint, H.; Löw, P.; Wallén, P.; Grillner, S. Cerebrospinal Fluid-Contacting Neurons Sense pH Changes and Motion in the Hypothalamus. J. Neurosci. 2018, 38 (35), 7713–7724. 10.1523/JNEUROSCI.3359-17.2018.

(32) Dapino, A.; Curti, S. Coincidence Detection Supported by Electrical Synapses Is Shaped by the D-Type K+ Current. J Gen Physiol 2026, 158 (2), e202513883. 10.1085/jgp.202513883.

(33) Rook, M. L.; Musgaard, M.; MacLean, D. M. Coupling Structure with Function in Acid-Sensing Ion Channels: Challenges in Pursuit of Proton Sensors. J Physiol 2021, 599 (2), 417–430. 10.1113/JP278707.

(34) Djenoune, L.; Khabou, H.; Joubert, F.; Quan, F. B.; Nunes Figueiredo, S.; Bodineau, L.; Del Bene, F.; Burcklé, C.; Tostivint, H.; Wyart, C. Investigation of Spinal Cerebrospinal Fluid-Contacting Neurons Expressing PKD2L1: Evidence for a Conserved System from Fish to Primates. Front. Neuroanat. 2014, 8. 10.3389/fnana.2014.00026.

(35) Bruni, J. E.; Reddy, K. Ependyma of the Central Canal of the Rat Spinal Cord: A Light and Transmission Electron Microscopic Study. J Anat 1987, 152, 55–70.

(36) Bjugn, R.; Haugland, H. K.; Flood, P. R. Ultrastructure of the Mouse Spinal Cord Ependyma. J Anat 1988, 160, 117–125.

(37) Prieto, D.; Rehermann, M. I.; Fabbiani, G.; Vitar, M.; Trujillo-Cenóz, O.; Falco, M. V.; Cúparo, M.; Trigo, F. F.; Russo, R. E. A Filopodia-Based Dendritic Mechanosensory Compartment in CSF-Contacting Neurons. bioRxiv April 10, 2026, p 2026.04.09.713694. 10.64898/2026.04.09.713694.

(38) Lyckenvik, T.; Forsberg, M.; Johansson, K.; Axelsson, M.; Zetterberg, H.; Blennow, K.; Illes, S.; Wasling, P.; Hanse, E. Ion Concentrations in CSF and Serum Are Differentially and Precisely Regulated. Brain Communications 2025, 7 (3), fcaf201. 10.1093/braincomms/fcaf201.

(39) Su, Q.; Hu, F.; Liu, Y.; Ge, X.; Mei, C.; Yu, S.; Shen, A.; Zhou, Q.; Yan, C.; Lei, J.; Zhang, Y.; Liu, X.; Wang, T. Cryo-EM Structure of the Polycystic Kidney Disease-like Channel PKD2L1. Nat Commun 2018, 9, 1192. 10.1038/s41467-018-03606-0.

(40) Zhou, X.; Li, M.; Su, D.; Jia, Q.; Li, H.; Li, X.; Yang, J. Cryo-EM Structures of the Human Endolysosomal TRPML3 Channel in Three Distinct States. Nat Struct Mol Biol 2017, 24 (12), 1146–1154. 10.1038/nsmb.3502.

(41) Brickley, S. G.; Cull-Candy, S. G.; Farrant, M. Development of a Tonic Form of Synaptic Inhibition in Rat Cerebellar Granule Cells Resulting from Persistent Activation of GABAA Receptors. J Physiol 1996, 497 *( Pt* *3*) (Pt 3), 753–759. 10.1113/jphysiol.1996.sp021806.

(42) Kawaguchi, H.; Yamanaka, A.; Uchida, K.; Shibasaki, K.; Sokabe, T.; Maruyama, Y.; Yanagawa, Y.; Murakami, S.; Tominaga, M. Activation of Polycystic Kidney Disease-2-like 1 (PKD2L1)-PKD1L3 Complex by Acid in Mouse Taste Cells. J Biol Chem 2010, 285 (23), 17277–17281. 10.1074/jbc.C110.132944.

(43) Prendergast, A. E.; Jim, K. K.; Marnas, H.; Desban, L.; Quan, F. B.; Djenoune, L.; Laghi, V.; Hocquemiller, A.; Lunsford, E. T.; Roussel, J.; Keiser, L.; Lejeune, F.-X.; Dhanasekar, M.; Bardet, P.-L.; Levraud, J.-P.; Beek, D. van de; Vandenbroucke-Grauls, C. M. J. E.; Wyart, C. CSF-Contacting Neurons Respond to Streptococcus Pneumoniae and Promote Host Survival during Central Nervous System Infection. Current Biology 2023, 33 (5), 940–956.e10. 10.1016/j.cub.2023.01.039.

(44) Bellegarda, C.; Zavard, G.; Moisan, L.; Brochard-Wyart, F.; Joanny, J.-F.; Gray, R. S.; Cantaut-Belarif, Y.; Wyart, C. The Reissner Fiber under Tension in Vivo Shows Dynamic Interaction with Ciliated Cells Contacting the Cerebrospinal Fluid. Elife 2023, 12, e86175. 10.7554/eLife.86175.

(45) Uchitel, O. D.; González Inchauspe, C.; Weissmann, C. Synaptic Signals Mediated by Protons and Acid-Sensing Ion Channels. Synapse 2019, 73 (10), e22120. 10.1002/syn.22120.

(46) Delling, M.; DeCaen, P. G.; Doerner, J. F.; Febvay, S.; Clapham, D. E. Primary Cilia Are Specialized Calcium Signaling Organelles. Nature 2013, 504 (7479), 311–314. 10.1038/nature12833.

(47) Cristofori-Armstrong, B.; Rash, L. D. Acid-Sensing Ion Channel (ASIC) Structure and Function: Insights from Spider, Snake and Sea Anemone Venoms. Neuropharmacology 2017, 127, 173–184. 10.1016/j.neuropharm.2017.04.042.

(48) Zhang, Y.; Muyrers, J. P.; Testa, G.; Stewart, A. F. DNA Cloning by Homologous Recombination in Escherichia Coli. Nat Biotechnol 2000, 18 (12), 1314–1317. 10.1038/82449.

(49) Hamill, O. P.; Marty, A.; Neher, E.; Sakmann, B.; Sigworth, F. J. Improved Patch-Clamp Techniques for High-Resolution Current Recording from Cells and Cell-Free Membrane Patches. Pflugers Arch. 1981, 391 (2), 85–100.

(50) Zorrilla de San Martin, J.; Jalil, A.; Trigo, F. F. Impact of Single-Site Axonal GABAergic Synaptic Events on Cerebellar Interneuron Activity. J. Gen. Physiol. 2015, 146 (6), 477–493. 10.1085/jgp.201511506.

(51) A first morphological and electrophysiological characterization of Fañanas cells of the mouse cerebellum - Singer - 2025 - The Journal of Physiology - Wiley Online Library. https://physoc.onlinelibrary.wiley.com/doi/10.1113/JP285949 (accessed 2026-05-04).

(52) Schindelin, J.; Arganda-Carreras, I.; Frise, E.; Kaynig, V.; Longair, M.; Pietzsch, T.; Preibisch, S.; Rueden, C.; Saalfeld, S.; Schmid, B.; Tinevez, J.-Y.; White, D. J.; Hartenstein, V.; Eliceiri, K.; Tomancak, P.; Cardona, A. Fiji: An Open-Source Platform for Biological-Image Analysis. Nat Methods 2012, 9 (7), 676–682. 10.1038/nmeth.2019.

(53) Rothman, J. S.; Silver, R. A. NeuroMatic: An Integrated Open-Source Software Toolkit for Acquisition, Analysis and Simulation of Electrophysiological Data. Front. Neuroinform. 2018, 12. 10.3389/fninf.2018.00014.

(54) Bright, D.; Smart, T. G. Methods for Recording and Measuring Tonic GABAA Receptor-Mediated Inhibition. Front. Neural Circuits 2013, 7. 10.3389/fncir.2013.00193.

(55) Gibbons, B. H.; Edsall, J. T. RATE OF HYDRATION OF CARBON DIOXIDE AND DEHYDRATION OF CARBONIC ACID AT 25 DEGREES. J Biol Chem 1963, 238, 3502–3507.

(56) Ho, C.; Sturtevant, J. M. THE KINETICS OF THE HYDRATION OF CARBON DIOXIDE AT 25 DEGREES. J Biol Chem 1963, 238, 3499–3501.

(57) Eigen, M. Proton Transfer, Acid-Base Catalysis, and Enzymatic Hydrolysis. Part I: ELEMENTARY PROCESSES. Angewandte Chemie International Edition in English 1964, 3 (1), 1–19. 10.1002/anie.196400011.

(58) Hu, M.; Liu, Y.; Wu, J.; Liu, X. Influx-Operated Ca2+ Entry via PKD2-L1 and PKD1-L3 Channels Facilitates Sensory Responses to Polymodal Transient Stimuli. Cell Reports 2015, 13 (4), 798–811. 10.1016/j.celrep.2015.09.041.

